# Population dynamics of the thalamic head direction system during drift and reorientation

**DOI:** 10.1101/2021.08.30.458266

**Authors:** Zaki Ajabi, Alexandra T. Keinath, Xue-Xin Wei, Mark P. Brandon

## Abstract

The head direction (HD) system is classically modeled as a ring attractor network^1,2^ which ensures a stable representation of the animal’s head direction. This unidimensional description popularized the view of the HD system as the brain’s internal compass^3,4^. However, unlike a globally consistent magnetic compass, the orientation of the HD system is dynamic, depends on local cues and exhibits remapping across familiar environments^5^. Such a system requires mechanisms to remember and align to familiar landmarks, which may not be well described within the classic 1-dimensional framework. To search for these mechanisms, we performed large population recordings of mouse thalamic HD cells using calcium imaging, during controlled manipulations of a visual landmark in a familiar environment. First, we find that realignment of the system was associated with a continuous rotation of the HD network representation. The speed and angular distance of this rotation was predicted by a 2^nd^ dimension to the ring attractor which we refer to as network gain, i.e. the instantaneous population firing rate. Moreover, the 360-degree azimuthal profile of network gain, during darkness, maintained a ‘memory trace’ of a previously displayed visual landmark. In a 2^nd^ experiment, brief presentations of a rotated landmark revealed an attraction of the network back to its initial orientation, suggesting a time-dependent mechanism underlying the formation of these network gain memory traces. Finally, in a 3^rd^ experiment, continuous rotation of a visual landmark induced a similar rotation of the HD representation which persisted following removal of the landmark, demonstrating that HD network orientation is subject to experience-dependent recalibration. Together, these results provide new mechanistic insights into how the neural compass flexibly adapts to environmental cues to maintain a reliable representation of the head direction.

## Main Text

The head direction (HD) system is commonly referred to as a neural compass, supporting a navigator’s sense of direction^3,4,6–9^. However, unlike a traditional compass, the orientation of the HD system is not globally consistent, but is instead anchored to local environmental cues^10–12^. While updating the internal HD representation requires integration of information from multiple streams (i.e., vestibular, motor, visual, etc…)^3,13–17^, manipulations of visual cues alone are sufficient to reorient this representation^18–22^. Thus, visual input often exerts a dominant influence on the HD network alignment, likely through a continuous feedback correction that calibrates the integration of angular movements and prevents the internal HD representation from drifting^23^. The interactions between the visual input and HD neurons are not fully understood, however, computational models of the HD network suggest that plasticity mediates the integration of visual information within the network^23–26^, which was experimentally confirmed, in fruit flies^27–29^. While these studies focus on the role of plasticity in the stabilization of the HD system, during navigation, when visual information is available, we do not know whether plastic effects could persist once the visual input is removed and whether they participate in stabilizing the internal HD estimation during darkness. Here, we characterize the thalamic HD network response to various visual manipulations in freely behaving mice, yielding novel insights into the dynamic mechanisms underlying anchoring and calibration of this representation, in light and dark conditions.

### Characterizing the HD network via calcium imaging

We obtained simultaneous recordings of up to 255 anterodorsal thalamic (ADN) cells simultaneously via a miniaturized head-mounted endoscope^30–32^ as mice freely explored a small elevated circular platform inside a larger enclosed chamber (Fig. 1a-g, Extended Data Fig. 1, Methods). This chamber was composed of a fully-encompassing 360° circular LED screen covered by a black plastic dome with a central oculus for behavioral camera access (Fig. 1a). During habituation and baseline recordings at the start of each session, we displayed a polarizing vertical white stripe. All subsequent testing involved the manipulation of this visual cue.

**Figure 1:**
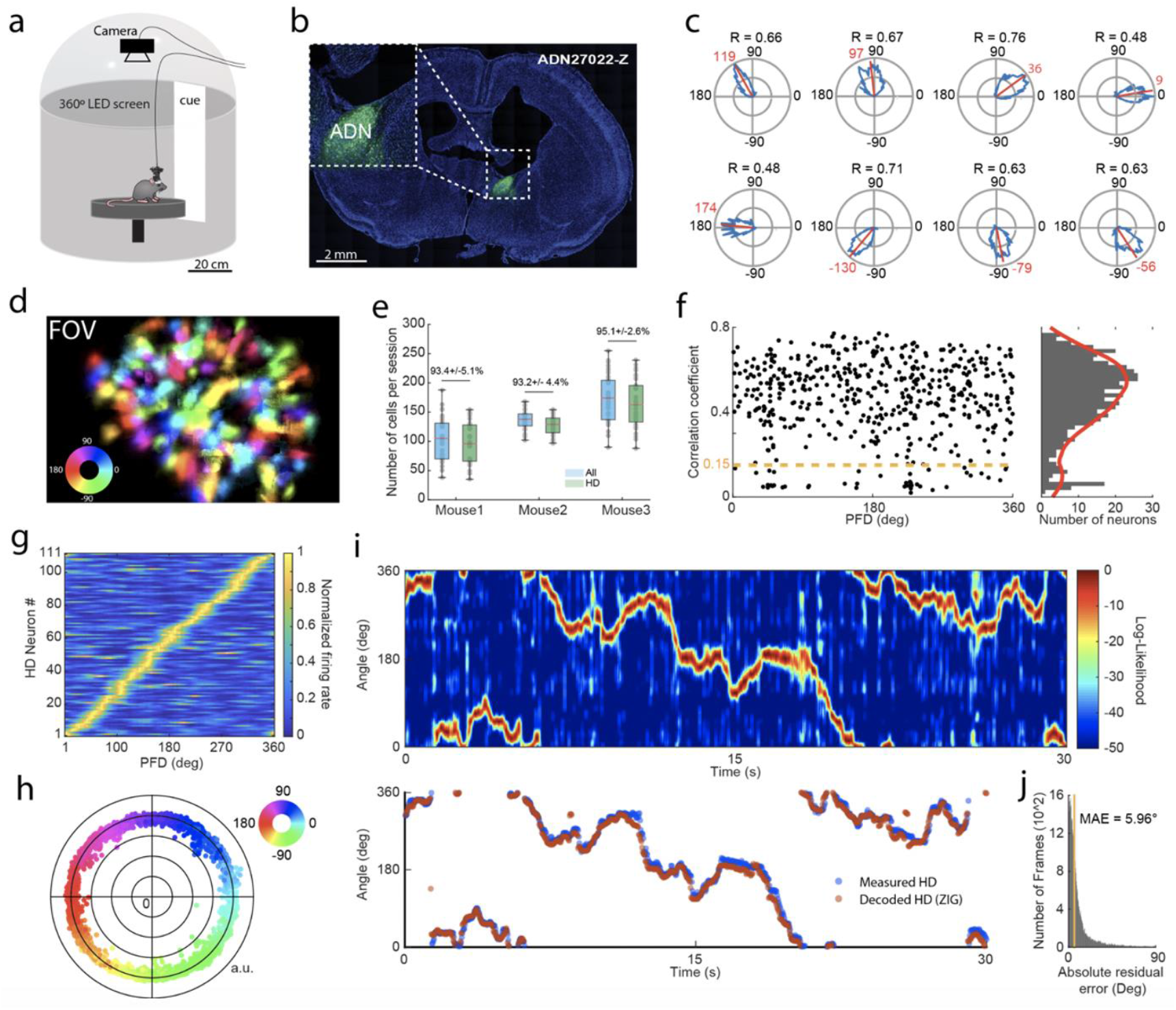
Population recordings in mouse ADN. **a**. Schematic showing recording environment with 360° LED screen. **b**. GCaMP6f expression in the ADN. **c**. Example tuning curves of ADN cells in polar coordinates. Red lines and numbers show the Mean Resultant Vectors (MRV) and preferred firing direction (PFD), respectively. R: Correlation coefficient. **d**. Field of view (FOV) of the ADN showing PFDs of each cell. **e**. Distribution of ADN cells recorded across mice (n = 3) and sessions (n = 99). Red line indicates median. Values above boxplots indicate percentage of HD cells (Green) among all recorded ADN cells (Blue) shown as ‘mean ± STD’. **f**. Example distribution of correlation coefficients of ADN cells. Dashed yellow line represents the HD-neuron detection threshold (Shuffled control: p < 0.05). Data shown from three baseline recording sessions of 10 minutes each (one per mouse). **g**. HD population coverage of the azimuthal plane from one recording session. **h**. Projection of high dimensional neural data onto a 2D polar plane using a feedforward neural network, during a baseline recording. **i**. Decoding head-direction from neural activity using the ZIG model. Top: Log-likelihood distribution across time. Bottom: measured HD (blue) and decoded HD (red) using maximum likelihood. **j**. Distribution of absolute residual error across 42 baseline epochs of three minutes each. MAE: median absolute error.

Calcium imaging data were motion-corrected and fluorescent transients of putative cells and spiking activity was inferred from extracted transients^33,34^ (Extended Data Fig. 2a). Baseline recordings revealed that simultaneously recorded HD cells exhibited preferred firing directions (PFDs) which tiled the full 360° of the horizontal plane (Fig. 1g, Extended Data Fig. 1c). Consistent with prior *in vivo* electrophysiological studies, the majority of ADN neurons were tuned to specific azimuthal head directions^3,35,36^ albeit with higher proportions (Fig. 1c-g, Extended Data Fig. 1), tuning was stable in the presence of visual cues^36^ (Extended Data Fig. 1), HD cells exhibited anticipatory firing^37^ (Extended Data Fig. 3a), and unlike the HD system in central complex of drosophila^38^ we did not observe topographic organization of HD cells by PFD (Fig. 1d, Extended Data Fig. 1b).

To infer the internal representation of HD from the activity of the HD-cell population (referred to, here, as the *internal HD*), we trained a zero-inflated-Gamma (ZIG) Bayesian decoder^39^ to estimate the animal’s HD on the basis of the baseline training data from each session (Fig. 1i). This decoder accurately recovered the mouse’s HD in stable experimental conditions (median absolute error on test data = 5.96°; Fig. 1i, j) and establish that this approach is well-suited to analyze the HD network from a large population perspective.

To visualize a 2^nd^ dimension of the representational space of the head direction network, we developed a method to project our HD population data into a 2-dimensional polar state-space (Methods). By imposing circularity on the first dimension, the second dimension captures variability in the neural data that cannot be explained by head direction. When applied to the baseline data, we obtain a ring-like structure (Fig. 1h), reminiscent of both attractor HD network models and prior analyses^1,40,41^.

### Network gain covaries with resetting dynamics during cue manipulation

From a theoretical standpoint, allowing states to occupy varying radii, in latent space, is akin to allowing the HD system to transition between energy states, assuming that the radius is a certain reflection of global activity in the network. We hypothesized that, when the same external inputs are applied to rotate the network, state transitions would be fastest at the lower end of the radial component because of the decreasing distance between states representing different angles, near the center of the baseline ring. Modulation of the state radius might thus provide the HD system with an efficient mechanism to rapidly reorient. To test this hypothesis, we first examined network responses to instantaneous cue removal and reappearance in new positions after darkness. Following a baseline recording, the cue was removed for two minutes (*darkness*) after which it reappeared at a 90° shifted position for two minutes. This sequence resulted in network drift during darkness and ‘reset’ events when the system reoriented to the visual cue. We repeated this sequence four times per recording session (Fig. 2a).

**Figure 2:**
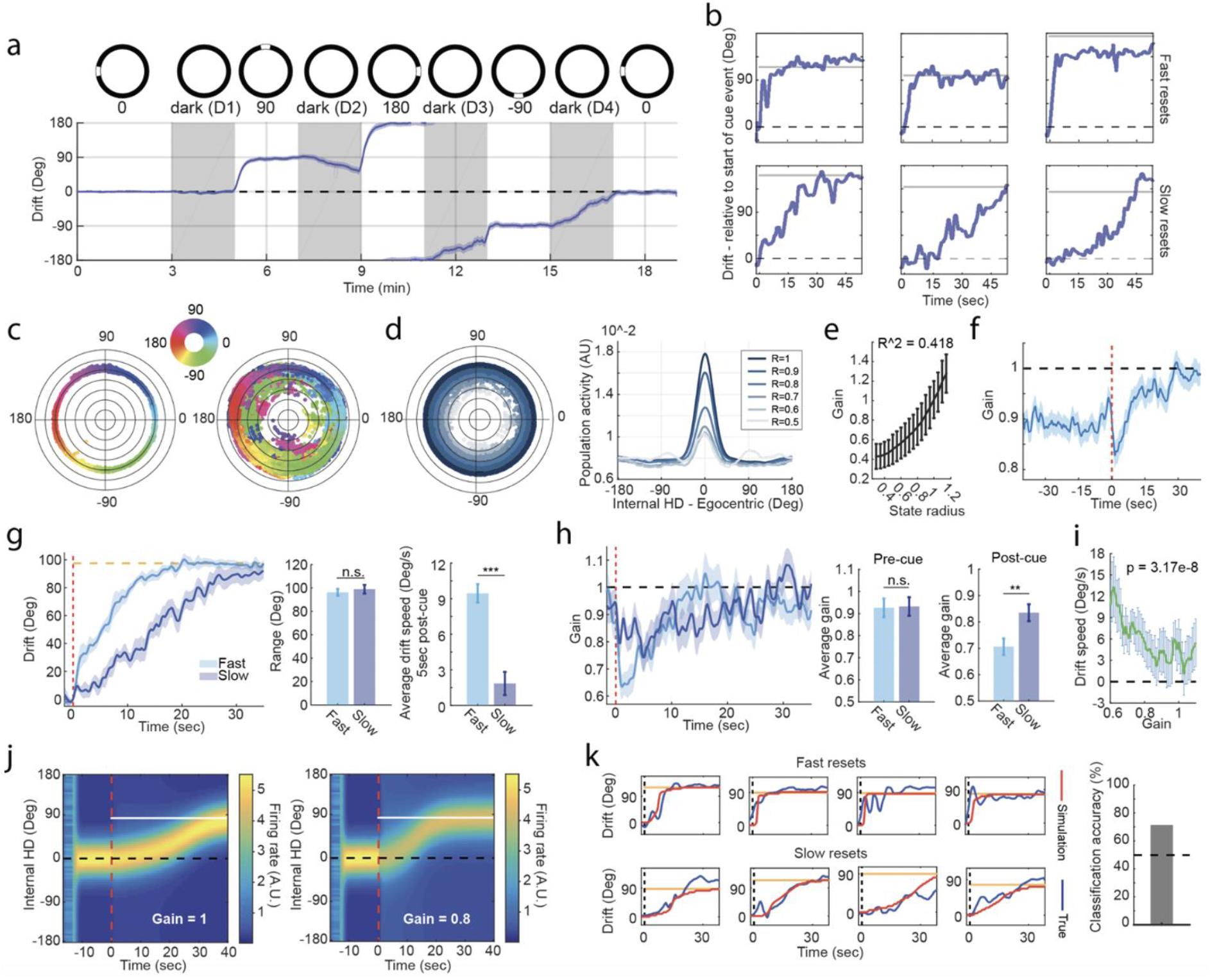
Network gain covaries with resetting dynamics. **a**. Mean population drift (from baseline cue condition) during the two-minute cue-shift experiment (n = 42 sessions). **b**. Examples of fast (top row) and slow (bottom row) resets. Horizontal solid line is the cue location. Drift values are relative to the drift angle at the moment of cue onset. **c**. Example projection of population activity onto a low dimensional 2D polar plane to highlight structural differences between baseline (left) and the entire session (right). **d**. State radius versus population activity. Left: same as in (**d**) however state points are color coded according to their radius. Right: Reconstructed mean bump of activity in egocentric reference frame across radius ranges. **e**. Relationship between network gain and state radius. The R-squared value corresponds to a linear regression model fit. Error bars indicate mean and STD. **f**. Triggered average of network gain (n= 168 = 4×42 cue events). Dotted red line indicates moment of cue-display. **g**. Mean drifts for fast (light blue; n = 22 resets) and slow (dark blue; n = 20 resets) resets. Dotted red line indicates moment of cue-display. While the two groups have similar ranges (Wilcoxon rank sum test: p = 0.4131, Z = 0.82), their speeds are different (Wilcoxon rank sum test: p = 1.0982e-6, Z = 4.87; 150 frames (~5s) post-cue). **h**. Network gains for the fast and slow reset-groups have similar amplitudes prior to cue-display (Wilcoxon rank sum test: p = 0.6234, Z = 0.49; 50 frames (~1.67s) pre-cue), yet they contrast significantly after cue-display (Wilcoxon rank sum test: p = 0.0085, Z = 2.63, RS = 535; 150 frames (~5s) post-cue). **i**. Relationship between gain and reset speed. Error bars indicate mean and SEM. **j**. Model simulation of the bump of activity during resets, showing gain control of reset speed. The gain remains constant (indicated value) after cue-display (dasher red line) for the rest of the simulation. Solid white line indicates relative cue location. **k**. Examples showing model-based prediction of reset (red) and true reset (blue). Dashed black line indicates moment of cue-display. Yellow line indicates cue location relative to drift at cue-display (71.43% classification accuracy). Time-dependent signals, in **a**, **f**, **g** and **h** are shown as mean (solid line) and SEM (shaded area) and bar graphs indicate mean and SEM

To characterize these network dynamics, we defined drift as the amount of mismatch between the measured HD and the decoded HD (Methods). Tracking this signal over the entire recording session, we find jumps following cue reappearance, which we will refer to as ‘resets’ (Fig. 2a). Notably, resetting events were not homogeneous: they occurred across a wide range of angles and at different speeds (Fig. 2b).

Cue manipulation also induced marked changes in the overall network activity, which coincided with changes in the radius of the latent space (Fig. 2c, d). To quantify this relationship, we computed the amount of population activity at each point in time relative to baseline, a measure we refer to as the *network gain*. State-space radius was highly correlated with network gain (Fig. 2d, e), indicating that gain can be used as an accessible and interpretable measure of the radial component. To better understand the relationship between resetting events and network gain, we first analyzed the 90°-centered reset range (i.e. [70:110]°-range). We found that the speed of rotation, or ‘reset speed’, was anticorrelated with network gain, consistent with our predictions (Fig. 2i). Grouping resetting events by their speed in the first five seconds post cue reappearance (Fig. 2g; Methods) revealed that fast resets were associated with a substantial reduction in gain, while slow resets exhibited a significantly smaller reduction in gain (Fig. 2h, Extended Data Fig. 4a). In all cases, resets took the form of a continuous rotation of the HD network from an initial orientation to the reset direction, passing by all intermediate angles with no visible appearance of secondary bumps in the population activity (Extended Data Fig. 5). This reset was surprisingly much slower than what has previously been estimated^21^, potentially due to differences in animal behavior, habituation to the environment and the circularly symmetric geometry of the experimental setup. Notably, an attractor network model that included gain modulation replicated these dynamics and reached 71.43% accuracy in classifying fast versus slow reset events from real data (Fig. 2j, k, Extended Data Fig. 6; Supplementary Material, *Attractor Network Model*).

Behavioral differences before and after cue onset were not significant and could not explain the sharp decrease in gain amplitudes (Extended Data Fig. 7a). However, reduced activity, as measured by the absolute head angular velocity, immediately preceding cue events, was predictive of fast resets, and vice versa (Extended Data Fig. 7b, c).

Resetting events also varied in the angular difference between their initial and stabilizing orientations. We grouped resets by the distance between pre-cue drift and its value after stabilization following cue reappearance (Extended Data Figs. 4b, 8a). We found that network gain was also anticorrelated with reset-range. As the amount of angular correction needed to realign the internal representation with the visual reference frame increased, the gain gradually dropped right after cue display (Extended Data Fig. 8b, c). This relationship was independent of reset speed (Extended Data Fig. 8a) which indicates that the network gain may also be modulated by the estimated degree of error between the internal representation and the changing location of the visual cue.

We also note that we detected a rapid spike in gain at cue-onset which was largest in short-range resets (Extended Data Fig. 9), which may reflect visual inputs to the system, but detailed investigation into this finding was limited by the temporal resolution of calcium imaging.

### Network gain maintains a trace of the visual cue during darkness

Prior work has demonstrated that the HD network drifts in the absence of polarizing visual cues^5,10,42^. In dark conditions, drifts increase with time and complexity of outward trajectories in a path integration task^43^. First, we replicated the classic result of increased drift during darkness. During all four darkness epochs (D1 to D4, respectively), we observed an increase in the variability of drift relative to baseline (Fig. 3a, b, Extended Data Fig. 10a). This increase in drift coincided with an abrupt drop in network gain upon cue removal which persisted for the duration of the darkness epoch (Fig. 3c). Surprisingly, changes in network gain were highly dependent on the internal HD. When the internal HD pointed toward the internal location of the visual cue (0°), the reduction in gain was minimal; deviations from this direction resulted in more pronounced gain decreases (Fig. 3d, e). In other words, the HD neurons that fired when the animal was facing the cue during baseline have higher average firing rates during darkness, following a reset. This pattern of gain tuning persists across all darkness periods, however, the contrast between peak and trough became larger with time (Extended Data Fig. 11). The animals’ behavior did not seem to affect this gain profile in any meaningful way other than the amplitude modulation causing the gain to increase with the absolute head angular-velocity (AV) (Fig. 3e, Extended Data Fig. 12), consistent with previous observations^38,44^. Furthermore, network gain fluctuations did not correlate with any measurable distortion to the drift-speed landscape within the same state-space (Head AV vs Internal HD) (Extended Data Fig. 12b), which maintained similar patterns to baseline (Extended Data Fig. 3b). This observation draws a clear distinction from the rapid representational shifts seen during resets and may point to a completely different mechanism linking network gain and drifts, in dark conditions. One interpretation of these results is that the HD network keeps a ‘memory trace’ of the visual input even after it is removed which might be used as an internal reference to guide behavior, during darkness. To our knowledge, this is the first evidence of such long-term experience-dependent preferential firing in the thalamic HD cells.

**Figure 3:**
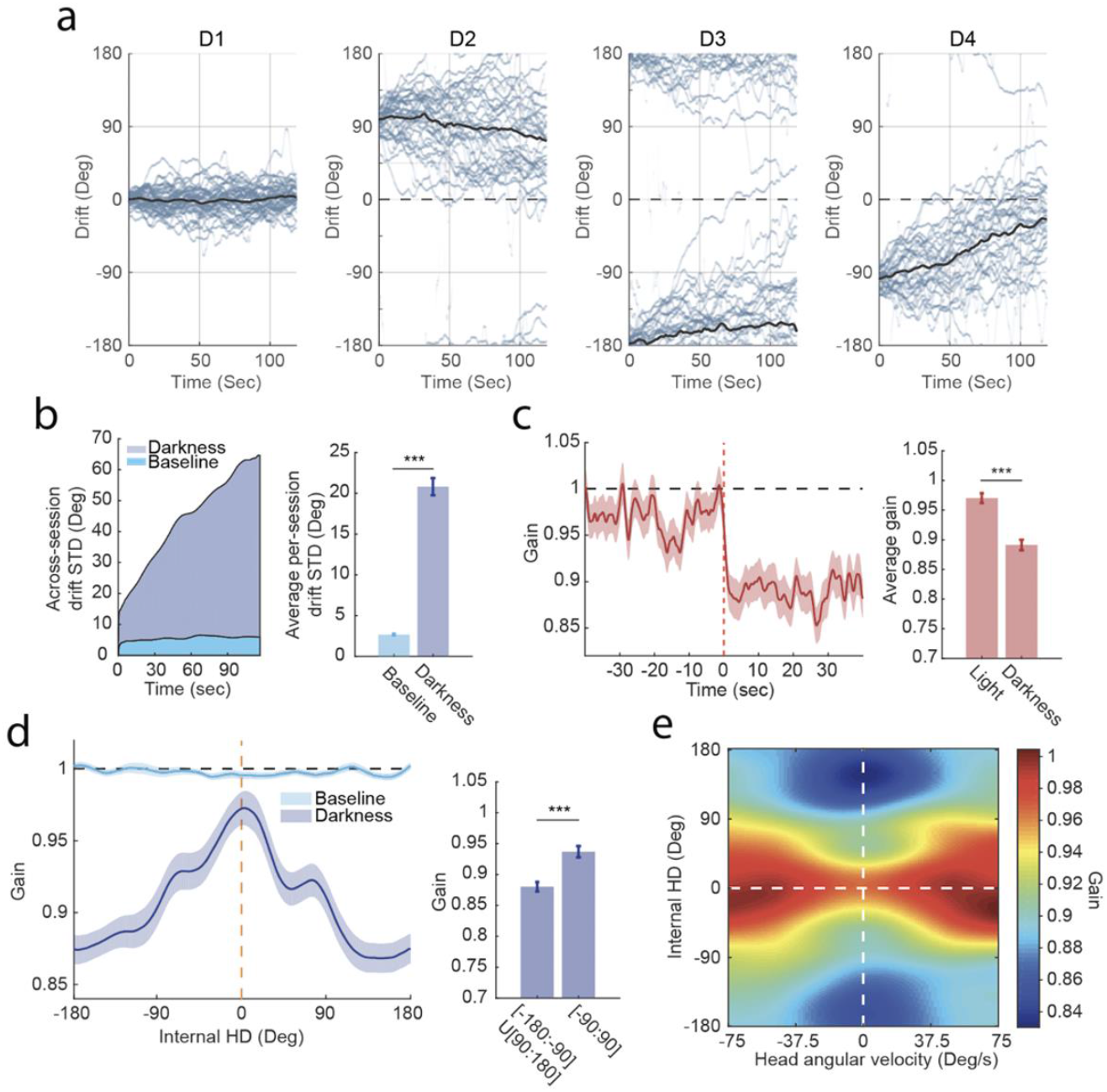
The network gain maintains a trace of the visual cue in darkness. **a**. All drift signals (light blue) across darkness periods D1 (n = 42), D2 (n = 35), D3 (n = 33) and D4 (n = 35), respectively. Black lines are mean drifts. For D2, D3 and D4, only darkness epochs that follow a correct reset are considered. **b**. Drift variability increases with time during dark sessions compared with baseline (baseline n = 42, darkness n = 145; Wilcoxon rank sum test: p = 7.5211e-23, Z = 9.84). **c**. Triggered-average of network gain shows an abrupt drop at the transition between cue-on and cue-off epochs, marked by the dotted red line (Wilcoxon rank sum test: p = 9.0165e-12, Z = 6.81, RS = 27362; comparison between means over 40 sec-pre and 40 sec-post cue-removal). **d**. Network gain tuning curves during BL (light blue) and darkness (dark blue). The internal HD is relative to the baseline cue location (dashed yellow line). The gain remains flat during baseline however, it peaks around the internal cue location ([−90:90]°) and drops sharply away from it ([−180:-90]U[90:180]°), in darkness (Wilcoxon rank sum test: p = 5.8683e-7, Z = 5.00). **e**. Network gain heatmap. Note the increase in amplitude and width of the gain tuning curve at larger head angular velocity (HAV). All CW sessions have been reflected across the x-axis and transformed into CCW ones. Signals in **c** and **d** are shown as mean (solid line) and SEM (shaded area) and bar graphs indicate mean and SEM

Although darkness induced an increase in drift during all four epochs, each epoch exhibited a different stereotyped pattern of drifts (Extended Data Fig. 10a, b). During D1, the HD representation became less stable than baseline. The drift trajectories fluctuated around the baseline orientation with no directional bias. Conversely, drift trajectories in D2, D3 and D4 exhibited substantial directional biases − albeit at relatively slow rates when compared to resets − dependent on the baseline orientation and prior cue locations. Specifically, during D2, drift tended to diverge from its reset orientation toward its baseline orientation, counter to the rotation implied by the previous cue shift. During D3 and D4, drift also tended to rotate toward the baseline orientation but consistent with the direction implied by the prior cue shifts. In comparison with D2, drifts were faster during D4 despite both conditions being preceded by symmetric cue rotations relative to baseline. These observations indicate that, during darkness following a reset (D2 to D4), the HD representation within the changing allocentric reference frame is less stable than baseline darkness (D1). Moreover, the persistent rotation of the reference frame, in one direction, appeared to bias drift in that direction (Extended Data Fig. 10c). Together, these results suggest that both the stable allocentric reference frame and the dynamic visual cue reference frame exert a persistent influence on the network orientation even after the visual cue is removed, possibly mediated by the dynamics of the network gain profile.

### Reference frame attraction in the HD network is time-dependent

Our results indicate that presentation of a rotated visual cue for two minutes was sufficient to cause a representational shift and override the influence of any available unaltered non-visual information (e.g. local olfactory cues, self-motion cues, etc). Yet, we observed a tendency of the network to rotate towards the initial baseline cue configuration, or *revert*, during darkness. To explore this and to better examine the potential influence of non-visual information linked to baseline representation, we performed an experiment in which we limited the display of the rotated visual cue to 20 seconds (alternating +/−90° from baseline, Fig. 4a). As in the prior experiment, these shortened cue events elicited resets followed by reversion towards baseline during darkness (Fig. 4b, c). However, in comparison with the second darkness epoch (D2) of the previous experiment - which was similarly preceded by a ±90° rotated cue event relative to baseline - we observed that reversion was much stronger following the 20s-visual-cue presentation (Fig. 4c). Vector field analysis was used to draw the dynamical landscape of the ‘drift-speed versus drift-angle’ state space, from observations (Methods). This allowed us to simulate drifts at various initial conditions which resulted in their convergence near baseline orientation, thus revealing the strong attraction of the baseline configuration (Fig. 4d). These results indicate that the internal representation of the baseline allocentric reference frame is not entirely lost after a reset and can still influence the HD network, in darkness, depending on the duration of experience within the competing reset reference frame context, which implicates plastic processes at play at some stage in this network.

**Figure 4:**
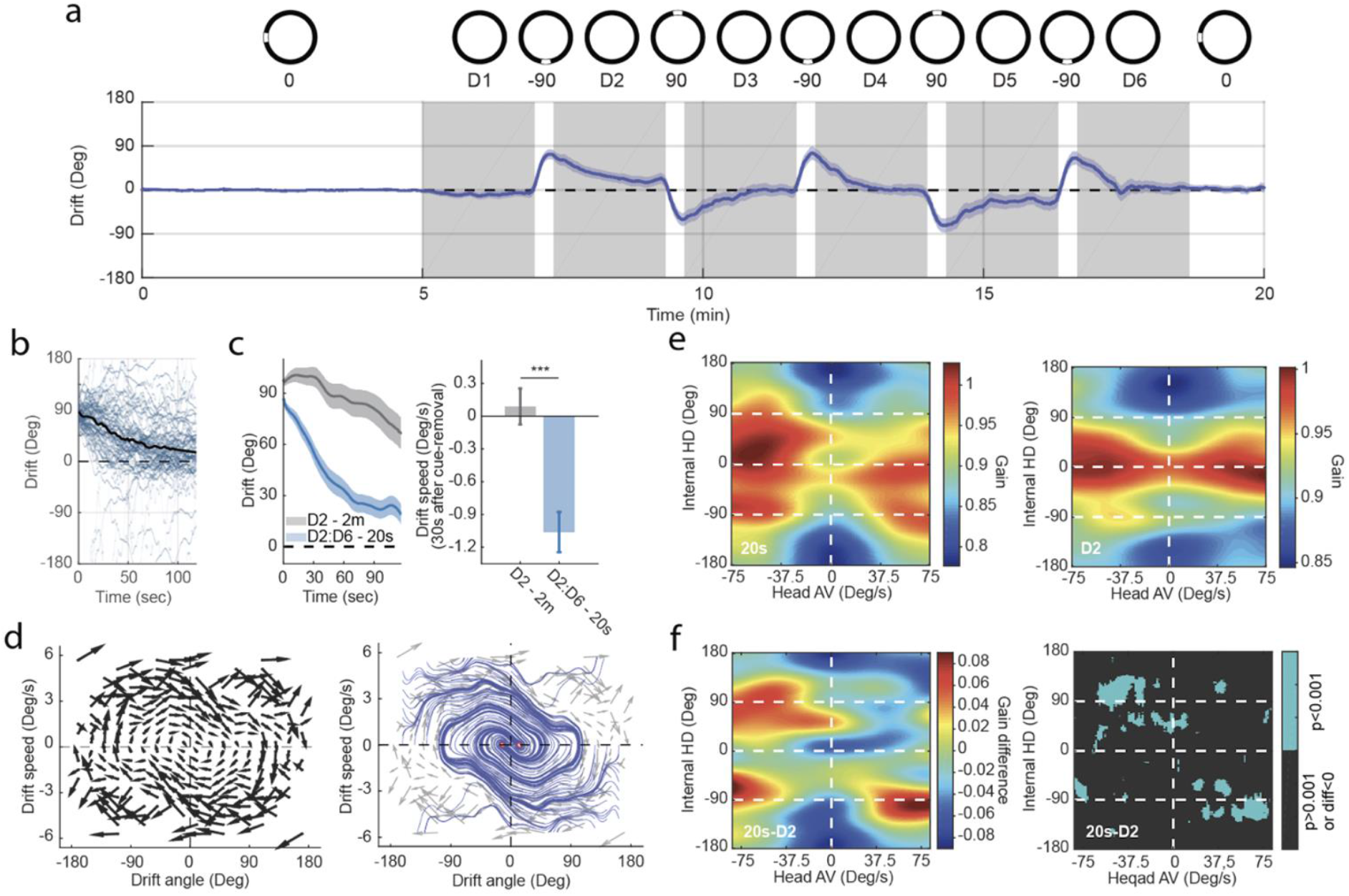
Attraction of internal representation to baseline reference frame is time dependent. **a**. Average population drift (from baseline) during the 20-second cue-shift experiment (n = 18 sessions). Darkness periods highlighted in grey. **b**. Individual drift signals following resets (light blue; n = 58) across darkness periods D2 to D6. Black line shows mean drift. Drifts following a −90° reset were reflected across the 0° axis. **c**. Mean drift signal during darkness in the 20s-cue-exposure experiment (dark blue) and in D2 of the 2m-cue-exposure experiment (grey). Mean drift-speed within the first 30s shows a strong reversion to baseline following a 20s cue-exposure (Wilcoxon rank sum test: p = 1.4264e-5, Z = 4.34). **d**. Left: Drift vector-field. Arrows point to the direction of mean drift-speed and mean drift-acceleration (n = 58). Arrows’ lengths were scaled down for illustration purposes. Right: Simulated streamlines over 1000 timesteps. The stable regime is highlighted in red. **e**. Network gain heatmaps. Left: 20s-cue-exposure experiment. data represents instances of reversion to baseline (n = 43). Right: D2 of the 2m-cue-exposure experiment (n = 35). **f**. Left: Gain difference (same data as in E) showing the appearance of new bumps at the locations of cue-shifts (±90°). Right: p-value matrix for data in left (Wilcoxon rank sum test; pixels where p>0.001 and/or gain(20s) < gain(D2) were marked as NaN). Time-dependent signals, in **a** and **c**, are shown as mean (solid line) and SEM (shaded area) and bar graph, in **c**, indicates mean and SEM

Next, we examined whether there was competition between reference frames, and how this would impact the network gain and drift dynamics. We began by characterizing gain as a function of internal HD and angular velocity. Whereas the gain profile during D2 exhibited a single peak around 0° in the internal HD (corresponding to the shifted cue orientation after reset), the gain profile during darkness following the 20s cue events exhibited additional peaks at ±90° (Fig. 4e, f). Notably, these peaks match the alternating ±90° cue structure of the experimental design, suggesting that gain profile differences reflected the experience with prior visual cues. In addition to the network gain profile, the drift pattern also showed systematic differences as a function of angular velocity and internal head direction between D2 and the darkness following 20s visual-cue display (Extended Data Figs. 13, 14). No obvious relationship between drift patterns and network gain profile could be determined, unlike what we observed during reset events, indicating that the relationship between gain and network state updating depends on the particular external input and/or current regime of the network.

### Dynamic visual cue updating induces a dynamic bias in the HD network

In the 2-minute experiment, visual information provided a dominant polarizing cue to reset the head-direction system. In some cases, resets were slow (>30 seconds) indicating that non-visual information competed with visual information to stabilize the network. In addition, the 20s experiment gave us further evidence that the baseline reference frame maintains a persistent influence on the HD network even after a cue-shift-induced reset. To better understand the dynamics of this competition, we tested whether visual information could drive resetting when in continuous conflict with all non-visual information, including self-motion cues, we recorded head direction cell populations during presentation of a slowly rotating visual cue (1.5 or 3.0°/s) for seven minutes (Fig. 5a, b). In all cases, and for both speeds, the head direction network was continuously updated by the rotating visual cue (Fig 5a-c) showing the dominant effect the visual input has over all other inputs in controlling the HD system. Unexpectedly, we found that, during darkness following the cue rotation, the HD network continued to rotate in the same direction and at a similar speed (Fig. 5a, b, d, Extended Data Figure 15), indicating that the rotating visual cue induced a persistent dynamic bias in the HD network which exerted an influence even in the absence of the visual input. This suggests that the visual flow can be used by the HD system to recalibrate the integration of angular velocity information (i.e. vestibular input) in order to anchor the internal HD representation to a dynamic visual reference frame. Moreover, we observed a similar attraction to the baseline internal representation as we saw in the 20s experiment. Overall, the system starts to stabilize once the internal HD representation comes close to realign with the initial reference frame (Fig. 5d), showing further evidence for the strong influence of the internal representation of the baseline reference frame and its potential role as an additional correction mechanism within the HD attractor network.

**Figure 5:**
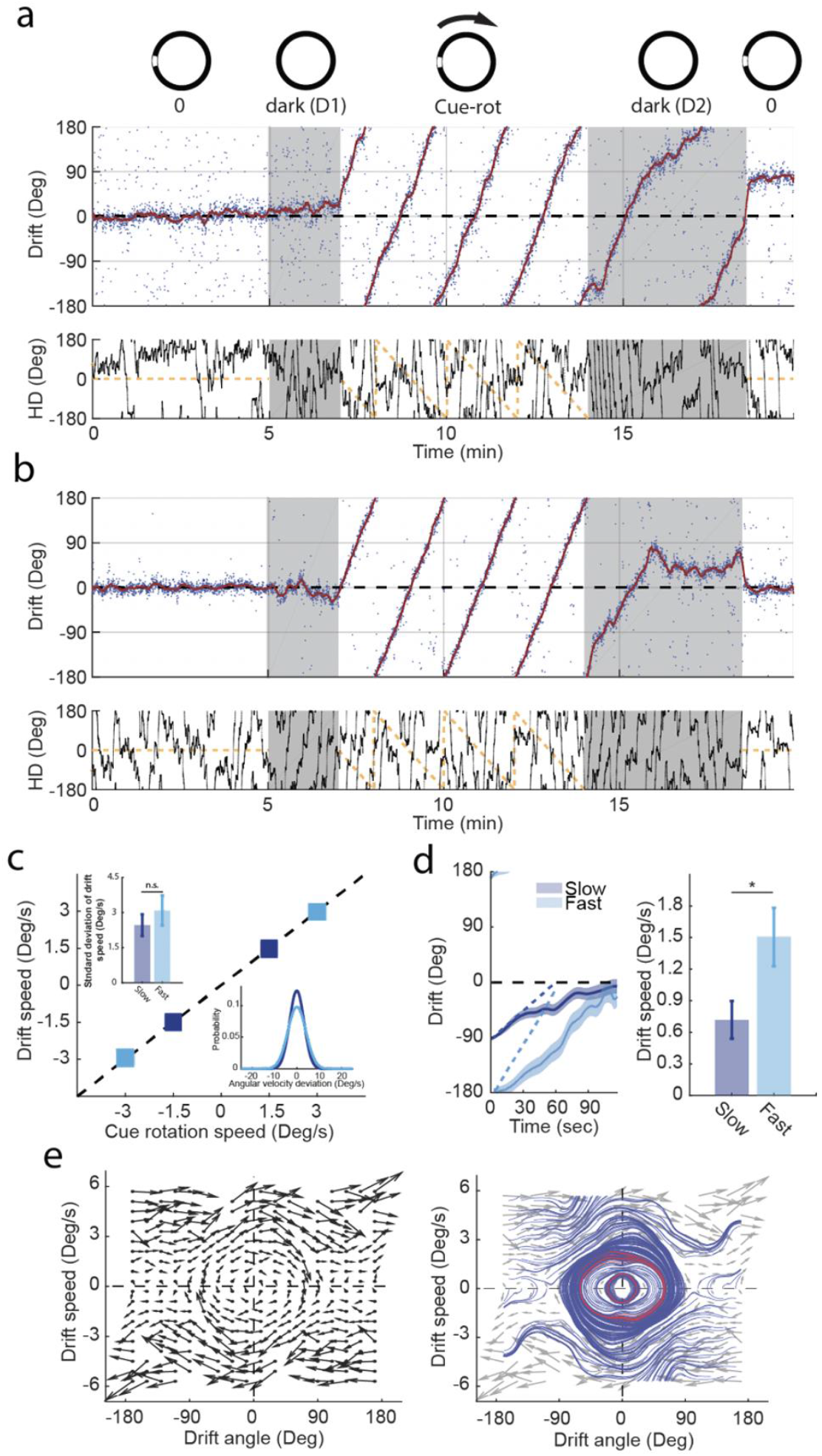
Optic flow calibrates head direction integration. **a**. Example population drift (from baseline) during fast cue-rotation (3°/s) experiment which exhibits persistent drift bias after cue removal (notice the remapping, at the end). Low-pass filtered (solid red) and unfiltered (dotted blue) drift signals are plotted on top of each other. Lower panel: Cue location (dashed yellow) and measured HD (solid black) relative to baseline cue-location. Darkness periods highlighted in grey. **b**. Example fast cue-rotation session showing stabilization of the internal representation with an overshoot past the baseline configuration. **c**. Mean drift-speed during cue-rotation for fast (light blue) and slow (dark blue) sessions. Error bars indicate across-session mean and STD. Top left insert: Comparison of drift speed STDs between fast and slow sessions (Wilcoxon rank sum test: p = 0.5262, Z = 0.63). Bottom right insert: Residual error between drift speed and actual cue angular-velocity. **d**. Left: Mean drift signal for fast (light blue; n = 19) and slow (dark blue; n = 25) sessions. Dotted lines correspond to the natural progression of the drift signal if the speed of drift matched the speed of the cue-rotation. Values are shown as mean (solid line) and SEM (shaded area). Right: Drift-speed comparison between fast and slow sessions within the first minute following cue-removal (Wilcoxon rank sum test: p=0.0393, Z=2.06,). Analysis was limited to the first 2 minutes because it includes sessions with 2-5 minutes of darkness post cue-rotation. **e**. Left: Drift vector-field. Arrows point to the direction of mean drift-speed and mean drift-acceleration (n = 60 sessions). Arrow lengths were scaled down for illustration purposes. Right: Simulated streamlines over 1000 timesteps. The stable regime is highlighted in red. In **a**, **b**, **c** and **d**, fast sessions where the drift angle, at the beginning of the 2nd darkness, is within [−180:−145]U[145:180]° were considered, while slow sessions where the drift’s initial position, in the 2nd darkness, is within [−125:−55]° were included. In **e**, all sessions were considered regardless of the drift angle at the beginning of the 2nd darkness. Bar graph, in **c** and **d**, indicates mean and SEM

## Discussion

Here we combined calcium imaging of large population recordings of ADN neurons in a visually controlled environment to examine how the mammalian HD network updates its representation. We show that controlled manipulations of a visual cue induce global fluctuations in network activity, which we term *network gain*. Network gain is not a simple product of ongoing sensory experience, but rather dynamically reflects the previous experience of the navigator: a polarizing visual landmark can induce lasting distortions in the network gain landscape even after the visual cue is removed. Furthermore, network gain is informatively linked to future network dynamics, as the reorientation of the HD representation following visual cue shifts can be predicted when network gain is integrated into a standard model of this system^45^. These results show that the network gain landscape can maintain a memory trace of a stable reference frame and suggest that this variable may mediate ongoing network dynamics. Finally, we show that the persistent influence of visual reference frames extends beyond the static gain modulation and can include induction of a persistent rotational bias following continuous rotation of the visual cue. Together, these results yield new insights into the dynamic influence of visual reference frames on the mammalian HD system.

The network gain drops that we observed during alignment of the HD network (i.e., reset) suggest the existence of a feedback control signal from brain regions downstream of ADN providing global inhibition to the network. A similar idea has been proposed in a study of the central complex of fruit flies^29,46^. The origin of such inhibition is yet to be discovered. Modulation of the global neural activity might allow the HD system to operate at different energy levels with varying degrees of stability, reflecting the level of confidence in the internal HD representation. We hypothesize that the animal’s engagement in exploratory behavior together with increased familiarity with the experimental environment and the specificity of the environmental geometry cause resistance to HD network reorientations imposed by visual cue shifts. This could explain the overall slower resets observed in our experiments when compared with previous reset studies^21^.

Recent work in fruit flies established a plastic relationship between the visual input and the compass neurons^27,28^, which are equivalent to HD neurons in the rodent brain. Their findings showed that the associations between compass neurons and visual scenes are time- and experience-dependent. The current work complements the previous studies and provides evidence for a time- and experience-dependent influence of visual landmarks, in mice. Indeed, the mammalian brain appears to maintain a memory of the associations between HD neurons and visual landmarks, in the form of preferential firing, long after the said landmarks disappear. We propose that memory traces of salient cues in ADN cells help stabilize the HD system during navigation, even in the absence of reliable environmental anchors such as in situations requiring path integration^43^.

Our results also indicate that memory traces from multiple reference frames can be found in the network gain landscape following short exposures to reset-inducing contexts (i.e. 20s cue-shifts). This apparent competition between conflicting reference frames results in predictable drifts of the HD network representation towards its most stable configuration, defined by the environmental context with the longer exposure. This attraction is likely achieved through integration of the unchanged non-visual information. We speculate that the underlying mechanisms leading to such behavior involve synaptic plasticity and that the 20s cue-events were insufficient to form new associations between HD neurons and the shifted visual reference frame. This may in turn explain the weaker reversions observed when the shifted-cue events were extended to two minutes which was long enough to promote new associations between the HD neurons and allocentric cues and enable a new stable state for the network, though future work is necessary to probe the mechanistic bases of these findings.

The fact that the persistent influence of a visual reference frame can act dynamically, as demonstrated in our cue rotation experiment, suggests that plasticity also acts at the level of integration of vestibular inputs and self-motion cues to allow for experience-dependent recalibration. Our findings complement a similar result discovered in place cells^23^ and support a model of hierarchical transfer of information from HD neurons to downstream cells of the navigation system (i.e., place cells, grid ells, etc.) in order to maintain consistent and flexible cognitive maps^3,29,47,48^.

Use of calcium imaging allowed us to obtain an order of magnitude increase in the number of thalamic head direction cells recorded^35^, ensuring accurate decoding of the animals’ azimuthal plane. One limitation of calcium imaging is its relatively slow dynamics, with rise and decay time constants of 10’s and 100’s of milliseconds, respectively, in response to neuronal spiking. As such, the firing rates of neurons must be inferred, and while our decoding results demonstrate the accuracy of this inference^33,39^, the use of electrophysiology would provide a direct indication of spiking activity. Although the development of linear probe technology remains promising^49^, the yield of neurons within small and deep nuclei, such as the ADN, remains a challenge for future development and experimentation.

Ultimately, our findings provide insights into the mechanisms that govern realignment and stabilization of the HD network, and how long-term effects of prior experience impact its dynamics. Importantly, these findings highlight some of the complexity of the internal HD representation and motivate studying this system in a multidimensional framework. The present work shows evidence for a functional interpretation of the global fluctuations in network activity (i.e., gain) when treated as a separate dimension. Future work, probing the origins of such fluctuations and allowing their perturbation will be critical to unveil the complete picture of the intrinsic structure of the HD representation.

## Methods

### Subjects

12 male wild-type mice (C57Bl/6, Charles River) were used for this study, three of which provided enough simultaneously recorded head direction cells for continued experimentation. Mice were housed individually on a 12-h light/dark cycle at 22 °C and 40% humidity with food and water ad libitum. All experiments were carried out in accordance with McGill University and Douglas Hospital Research Centre Animal Use and Care Committee (protocol #2015-7725) and in accordance with Canadian Institutes of Health Research guidelines.

### Surgeries

During all surgeries, mice were anesthetized via inhalation of a combination of oxygen and 5% Isoflurane before being transferred to the stereotaxic frame (David Kopf Instruments), where anesthesia was maintained via inhalation of oxygen and 0.5–2.5% Isoflurane for the duration of the surgery. Body temperature was maintained with a heating pad and eyes were hydrated with gel (Optixcare). Carprofen (10 ml kg−1) and saline (0.5 ml) were administered subcutaneously, respectively at the beginning and end of each surgery. Preparation for recordings involved three surgeries per mouse. First, at the age of seven to eight weeks, each mouse was injected with 600 nl of the non-diluted viral vector AAV9.syn.GCaMP6f.WPRE.eYFP, sourced from University of Pennsylvania Vector Core. All injections were administered via glass pipettes connected to a Nanoject II (Drummond Scientific) injector at a flow rate of 23 nl s−1. One week post-injection, a 0.5 mm diameter gradient refractive index (GRIN) relay lens (Go!Foton) was implanted above ADN (AP:1.8, ML:0.8, DV:−3). No aspiration was required. In addition to the GRIN lens, three stainless steel screws were threaded into the skull to stabilize the implant. Dental cement (C&B Metabond) was applied to secure the GRIN lens and anchor screws to the skull. A silicone adhesive (Kwik-Sil, World Precision Instruments) was applied to protect the top surface of the GRIN lens until the next surgery. Two weeks after lens implantation, an aluminum baseplate was affixed via dental cement (C&B Metabond) to the skull of the mouse, which would later secure the miniaturized fluorescent endoscope (miniscope) in place during recording. The miniscope/baseplate was mounted to a stereotaxic arm for lowering above the implanted GRIN lens until the field of view contained visible cell segments and dental cement was applied to affix the baseplate to the skull. A polyoxymethylene cap with a metal nut weighing ~3 g was affixed to the baseplate when the mice were not being recorded, to protect the baseplate and lens, as well as to simulate the weight of the miniscope. After surgery, animals were continuously monitored until they recovered. For the initial three days after surgery mice were provided with a soft diet supplemented with Carprofen for pain management (MediGel CPF). Screening and habituation to recording in the experimental environment began 2-3 days following the baseplate surgery. The first 3-4 weeks of recordings were used to confirm quality and reliability of the calcium data while the animal was exploring the environment with different screen displays.

### Data acquisition

In vivo calcium videos were recorded with a miniscope (v1; miniscope.org) containing a monochrome CMOS imaging sensor (MT9V032C12STM, ON Semiconductor) connected to a custom data acquisition (DAQ) box (miniscope.org) with a lightweight, flexible coaxial cable. The DAQ was connected to a PC with a USB 3.0 SuperSpeed cable and controlled with Miniscope custom acquisition software (miniscope.org). The outgoing excitation LED was set to 3–6%, depending on the mouse to maximize signal quality with the minimum possible excitation light to mitigate the risk of photobleaching. Gain was adjusted to match the dynamic range of the recorded video to the fluctuations of the calcium signal for each recording to avoid saturation. Behavioral video data were recorded by a webcam mounted above the environment. The DAQ simultaneously acquired behavioral and cellular imaging streams at 30 Hz as uncompressed avi files and all recorded frames were timestamped for post-hoc alignment. Two controllable LEDs (green and red) were added and used for tracking such that whenever the miniscope was attached to the baseplate, the green LED pointed to the right side of the mouse’s head and the red LED pointed to the left side. All other light sources from the miniscope were covered. All recordings took place inside a 360° LED screen (height: 1m, diameter: 90cm; Shenzhen Apexls Optoelectronic Co.) at the center of which we placed a wall-less circular platform (diameter: 20cm) raised 50cm above the ground. Mouse bedding was evenly spread over the platform before each recording session. In all recordings, mice were free to move on top of the raised platform. A half spherical dome was used to cover the environment and prevent external light from entering, while it also held the behavioral camera. The experimental environment was designed to maximize circular symmetry, in the absence of any screen display. During habituation, mice were recorded while exposed to a single vertical stripe or no visual display (darkness). These recordings were also used to confirm the quality of tracking the head-direction and the cue location, in different conditions. In all experiments of the present work, the ‘visual cue’ refers to a single white vertical stripe (Width: 15cm; Height: 1m).

### Data preprocessing

Calcium imaging data were preprocessed prior to analyses via a pipeline of open source MATLAB (MathWorks; version R2015a) functions to correct for motion artifacts^50^, segment cells and extract transients^34,51^ and infer the likelihood of spiking events via deconvolution of the transient trace through a first-order autoregressive model^33^. We wrote a Matlab (MathWorks, version 2015a) program to perform offline tracking of the LEDs and determine, at each frame, the animal’s head-direction. Another custom-written program was used to estimate the location of the visual. Both scripts were incorporated into the preprocessing pipeline.

### Data analysis

In this work, neural activity refers to the deconvolved calcium traces as described in (Friedrich et al., 2017)^33^ unless specified. The resulting time series (per neuron, per session) correspond to the inferred likelihood of spiking events. A moving average filter of width 3 frames (~100ms) is then applied on each time series. We refer to the obtained signal as ‘firing rate’.

### Identification of HD cells

For every identified cell segment (ROI), we construct a HD tuning curve by measuring the occupancy-normalized firing rate within each angle bin (1°/bin) of the horizontal plane (x-axis). The tuning curve is circularly smoothed with a moving average filter of width 50°. This allows us to have a better estimate of the angle bin that corresponds to the maximum firing rate of a given neuron’s tuning curve, which we will refer to as the preferred firing direction (PFD). Next, we construct a stimulus signal for that specific PFD by convolving the measured HD signal (from the behavioral camera) with a narrow Gaussian kernel (mean=PFD, std=17°) such that for every neuron *i*:

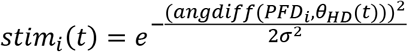

Where, *θ_HD_* is the measured HD time series, *σ* is the standard deviation of the Gaussian kernel and, *angdiff*(*a*, *b*) is a Matlab function that gives the subtraction of *a* from *b*, wrapped on the [−*π*, *π*] interval. We correlate the stimulus signal with a normalized version of the firing rate to obtain the Pearson correlation coefficient ‘*r*’ of each neuron. To determine the threshold value of *r* above which a cell can be identified as a HD neuron, we used data from 10 baseline recordings (3 minutes) per animal, randomly selected from the reset experiment. We start with a relatively high value *r_thresh_* and select all neurons such that *r* > *r_thresh_*. For each neuron, we produce 1000 shuffles of the firing rate using Matlab’s *circshift* function (to preserve the temporal correlation of the firing rate signal), at random shifts. We then correlate each shuffled version with the stimulus signal of the corresponding neuron in order to obtain a distribution of correlation coefficients (3 separate distributions, 1 per mouse). We define 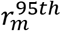 as the value that corresponds to the 95^th^ percentiles of the distribution, for mouse ‘m’. If 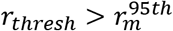, we keep iterating the same procedure while decreasing *r_thresh_* by 0.01 until convergence (i.e. 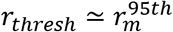) which constitutes the correlation coefficient threshold to identify HD neurons for mouse ‘m’ (see Extended Data Fig. 2 for illustration of the results).

### HD decoding from neural data

We trained a recently developed Bayesian decoder^39^ to decode the HD direction from the deconvolved calcium responses of the imaged neural population. Noise independence across neurons was assumed. Conceptually this decoder is similar to Bayesian decoding method for spike trains as commonly used in the literature^52^, except that we used zero-inflated-Gamma (ZIG) distribution to model the stochasticity of the deconvolved calcium responses, instead of Poisson distribution. Our previous results showed that the ZIG model could better capture the noise of the calcium signal and provide better decoding results compared to the Poisson noise model and a few other alternatives. Details of this procedure can be found in Section 4 of Wei et al. NBDT^39^. Here we smoothed the log-likelihood matrix (rows: angle bins, columns: frames) by iteratively summing the likelihoods over 5 frames (~166.7ms) centered around the corresponding timestep of each iteration, for each angle bin.

### Analysis of drift

We define drift as the difference between the measured head-direction (*θ_measured_*) and the decoded head-direction (*θ_decoded_*):

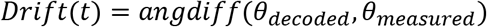

In all analyses involving drift calculation, both measured and decoded HDs were smoothed with a moving average filter of width 20 frames (~667ms). For the analysis of drift during darkness (except for heatmaps), further smoothing was applied to extract the low-frequency component of the signal whereby a moving average filter of width 300 frames (~10s) was used. In all cases, a simple linear regression was performed on the unwrapped drift signal over a sliding window of 20 frames (~667ms) to estimate the drift speed at the center of the regression window (i.e. slope of the fitted line).

### Separation of fast and slow resets

Classification of resets within the [70: 110]° range was done using Matlab’s *k-means* clustering function over the first 1450 frames following cue display. The algorithm separates between two clusters by generating 50 replicates with different initial cluster centroid positions for each replicate and then calculating the sums of point-to-centroid distances for each cluster using ‘cityblock’ as a distance metric.

### Reconstruction of the bump of activity

At any given time, we can reconstruct the bump of activity from the firing rates of each neuron and their respective tuning curves using a normalized weighted sum of tuning curves^53^:

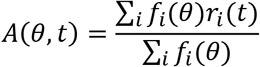

Where, A is a 360xT matrix (each row is a 1°-bin of the horizontal plane and each column is a frame within range T of the analysis), *f_i_* is the tuning curve of neuron *i* and, *r_i_* is the instantaneous firing rate of neuron *i*.

### Calculation of network gain

We assume that, at any given time, the thalamic HD network is subject to a global gain modulation of the firing rates, applied homogeneously on all ADN neurons such that:

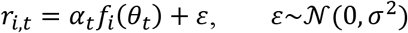

Where:

*r_i,t_*: instantaneous firing rate of ADN neuron *i*
*α_t_*: instantaneous gain factor
*f_i_*: tuning curve of ADN neuron *i*(determined from baseline)
*θ_t_*: Decoded head-direction (from neural activity), at time *t*
*ε*: Additive white Gaussian noise with standard deviation *σ*.

Our goal is to estimate at any given time *t*, the value of *α_t_*using maximum likelihood estimation (MLE) approach.

Given the decoded head-direction, at time *t*, *θ_t_*as well as the tuning curves *f_i_*for all ADN neurons, we obtain the likelihood of observing *r_i,t_*with parameter *α_t_*:

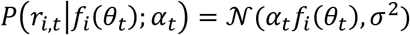

We define the vectors:

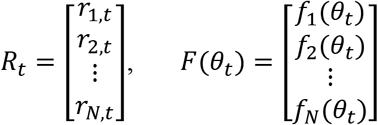

Where, *N*is the number of ADN neurons, in the network.

Assuming independent activity between said neurons, we can calculate the likelihood of observing *R_t_*:

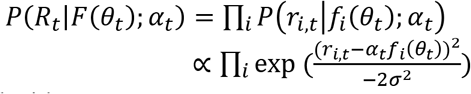

We apply the logarithm on both sides:

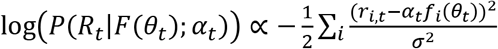

Our objective is to determine the parameter 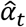that maximizes the log-likelihood such that:

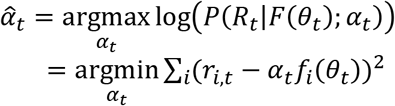

We take the derivative of the objective function w.r.t *α_t_*and set it to zero:

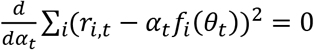

Thus:

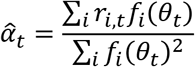

### HD network simulation

We designed an artificial neural network to simulate the behavior we see in the HD system. The network can maintain a stable HD representation (bump of activity) via lateral inhibition and input from the vestibular system (angular velocity (AV) cells). We incorporated gain modulation to allow for an internal control of drift speed, in reset situations. Details of the network architecture, neural dynamics and parameter optimization are detailed in additional supplementary material called *Attractor Network Model*.

### Gain heatmap analysis

Gain heatmaps are 2D matrices where each pixel *p*(*x*, *y*) is a 2D bin of width 1.5°/s corresponding to the measured angular head velocity and, height 1° corresponding to the decoded HD. Pixel *p*(*x*, *y*) represents the mean network gain – across mice and across sessions – within a 2D average window of width [x-3:x+3]°/s and height [y-15:y+15]°. A 2D Gaussian filter of standard deviation =15 (15° x 22.5°/s) is then applied. The network gain, the decoded HD and the measured HD were all smoothed with a moving average filter of width 20 frames (~667ms) while the measured head angular velocity was approximated by a simple linear regression with a regression window of similar width. To evaluate the significance of the difference between gain heatmaps (Fig.4G), we performed a Wilcoxon rank sum test to compare, at each pixel, the gain distributions within the 2D window of width [x-3:x+3]°/s and height [y-15:y+15]° between darkness epochs of the 20s experiment and D2 of the 2-minute experiment. As we are only interested in the significance of the positive values (indicating the appearance of new bumps), negative values as well as p-values>0.001 were marked as NaN (“Not a Number”).

### Drift-speed heatmap analysis

Drift-speed heatmaps were generated following the same approach as for gain heatmaps. However, drift speed was approximated by a simple linear regression with a regression window of width 20 frames (~667ms). The p-value matrix for drift speed difference between the 20s experiment and D2 of the 2-minute experiment was calculated as described above. However, only p-values>0.001 were marked as NaN.

### Vector field analysis

The purpose of this analysis is to illustrate baseline attractiveness. We define the state space (y-axis: Drift-speed (°/s); x-axis: Drift-angle (°)). We construct a vector field matrix by dividing the x-axis into 18 bins of width 20° each within the range [-180:180]°, and the y-axis into 20 bins of width 0.03°/s each, within the range [−3:3]°/s. At each bin (x,y), we calculate the mean drift-speed and mean drift acceleration, across mice and across sessions. The two latter quantities represent the velocity components (u,v) that determine the length and direction of the velocity vector. We assume the vector field has a central symmetry w.r.t the baseline point (0,0) because of the symmetry in the experimental design. So, we generate an image of the original vector field that is its reflection across the origin. The two versions are then averaged to produce the final 2D vector field. Streamlines are generated with Matlab’s ‘*streamline*’ function.

### Dimensionality reduction using feedforward neural network

We designed a deep artificial neural network that maps the high-dimensional input (neural) data onto the 2D polar space (radial and angular components). It is generally believed that the main function of the HD system is to provide an estimate of the HD at any given time. Since most studies of this network (including ours) are conducted while recording the neural activity in animals placed on horizontal planes, it is fair to assume that most of the variability in the activity of HD neural population can be captured by a single variable representing the angle faced by the animal, at an instant *t*, w.r.t a given allocentric reference frame. Indeed, previous works have shown that, in stable conditions, different dimensionality reduction methods^40,41^ would produce a circular manifold that can be fairly approximated in a unidimensional polar state-space with a fixed radius. Nevertheless, Chaudhuri et al. (2019)^40^ observed that the structure becomes more complex during slow-wave sleep (SWS). Our guiding hypothesis is that the intrinsic geometric structure of the neural activity in the HD network lies in a multidimensional state-space and that latent variables other than the angular component are needed to explain the variability in spiking data, during non-stable conditions such as resets and drift situations. Here we propose the simplest augmentation to the latent structure by adding a radial component that we expect to indicate instantaneous changes in global energy levels of the HD network. While we believe the true intrinsic dimensionality of the HD neural data is higher than two, the current paper mainly focuses on the necessity of at least a second dimension of the HD system during instability.

To test our hypothesis, we developed a deep artificial neural network to extract a secondary dimension while imposing circularity on the first one. Effectively, our method projects the high dimensional neural data onto the two dimensions of the polar space (angular dimension *θ* and radial dimension *R*). The radial component *R* is a latent variable that can take any non-negative value. We used a feedforward neural network with three parallel branches. Two of these branches have three fully connected hidden layers (referred to as ‘first’ and ‘second’ or, respectively, ‘*B*_1_’ and ‘*B*_2_’), while the third branch has two fully connected hidden layers (referred to as ‘middle’ or ‘*B_m_*’) (Extended Data Fig. 12). The input layer receives a Nx1 vector of neural activity from N ADN neurons, at time t (both calcium traces as well as firing rates from deconvolved spikes can be fed to the model). The output layer is composed of two units that are the results of multiplying the output *g_t_* of the middle branch with the output *z_1,t_* of the first branch, on one hand, and the output *z_2,t_* of the second branch, on the other hand, as illustrated in the diagram of Extended Fig. 16.

We train our model on baseline data. The objective is to find the set of weights *W* that minimize the distance between the network output 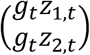 and the vector 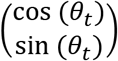. Where *θ_t_* is the measured head-direction of the animal at instant *t*. We define the loss function as the mean squared error:

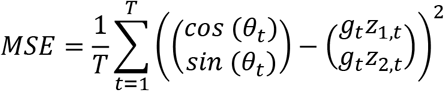

Where, *T* is the duration of the training epoch. If the algorithm converges, we obtain the following approximations:

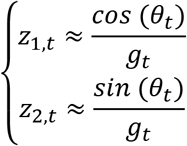

Let 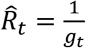, then we can rewrite the output of each branch:

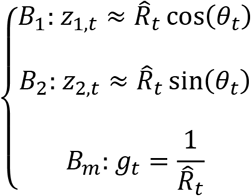

In effect, this would allow branches *B*_1_ and *B*_2_ to learn a mapping from the input (neural) space to the Cartesian transformation of the polar coordinates of a given state *s_t_*, at any time *t* (respectively, *B*_1_ projects the input onto the *x* axis and, *B*_2_ projects the input onto the *y* axis). While branch *B_m_* would learn a mapping from the input space to the inverse of the approximate radius 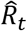 of said state, in polar space. If we assume 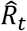 is a certain reflection of global neural activity, as per our hypothesis, then we expect small fluctuations of population activity, in the training data (baseline), to be sufficient to allow the network to extrapolate 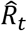 on test data with larger fluctuations.

### Statistics and reproducibility

All statistical tests are noted where the corresponding results are reported throughout the main text and supplement. All tests were uncorrected 2-tailed tests unless otherwise noted. Outliers were identified as data points that fall outside the mean±(3*std) range.

## Data availability

The complete dataset for all experiments is available upon request to the corresponding authors.

## Code availability

All source codes used in the current study are available upon request to the corresponding authors.

## Acknowledgements

We are grateful to S. Kim, H. C. Yong and A. Nieto-Posadas for the technical assistance and help gathering the histology data. We kindly thank L. Paninski, M. Hasselmo, S. Badrinarayanan, J. Ying, M. Yaghoubi and A. Peyrache for comments on previous versions of this paper. We also thank D. Aharoni for advice and assistance with the UCLA the UCLA miniscope. We are grateful to members of the Brandon laboratory for useful discussions, in particular M. Yaghoubi and R.R. Rozeske. This work was supported by funding from the Canadian Institutes of Health Research (Project grants #367017 and #377074), the Natural Sciences and Engineering and Research Council of Canada (Discovery grant #74105), and the Canada Research Chairs Program to M.P.B.

## Contributions

Z.A., A.T.K. and M.P.B. contributed to the experimental design; Z.A. performed all surgeries, recordings, data analysis, and modelling; X.X.W., A.T.K. and M.P.B. provided guidance for data analysis; Z.A. wrote the initial draft; All authors contributed to editing and revising the paper.

## Extended Data Figures

**Extended Data Figure 1.**
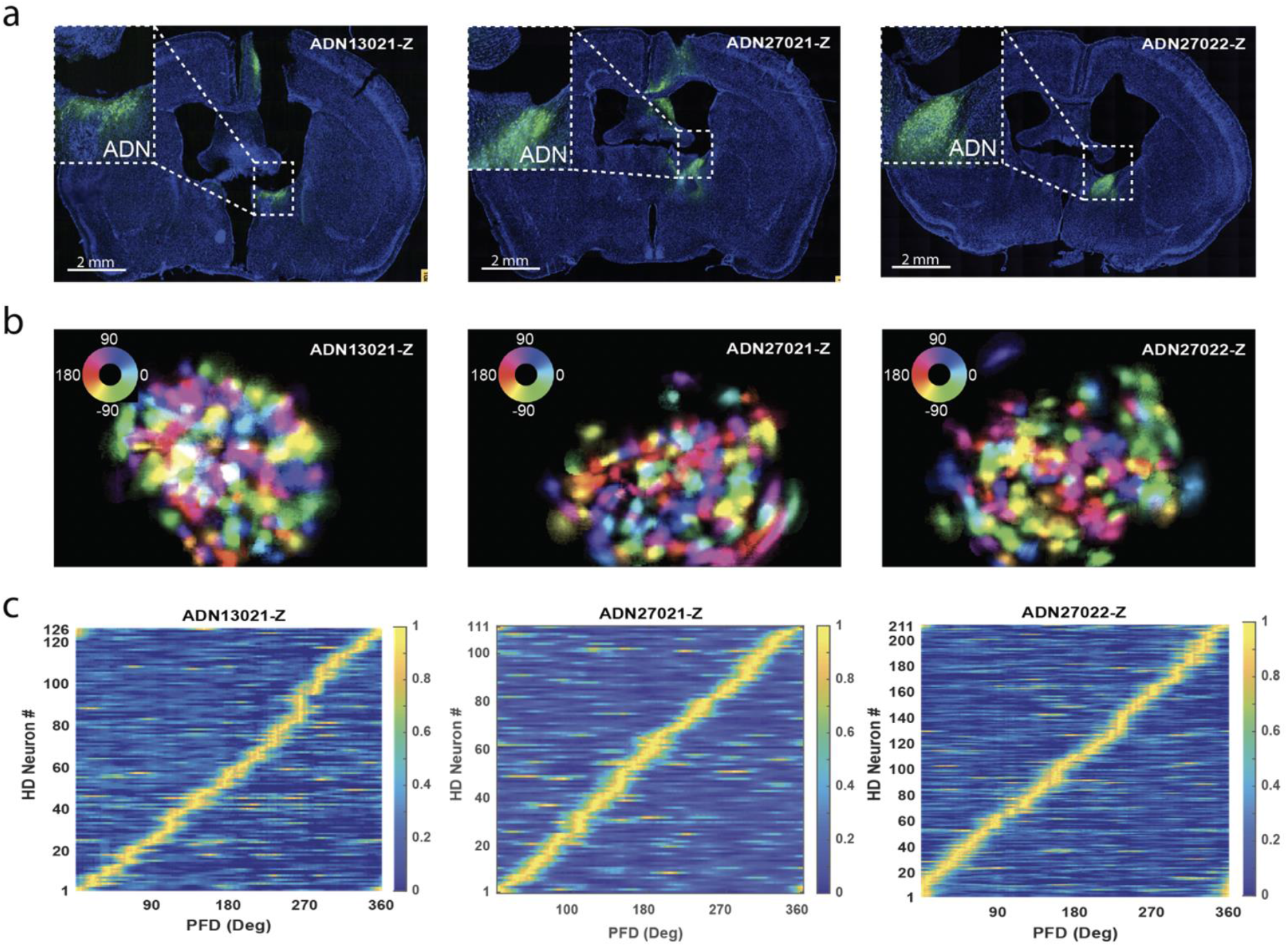
HD map of the anterodorsal thalamic nucleus (ADN). **a**. Histology data showing coronal brain sections from each mouse with GCaMP6f expression, in ADN region (anterior part). **b**. Directional maps of the ADN in each mouse. HD cells are colored according to their preferred firing direction (PFD). Color-wheel shows angle-color assignments. **c**. Examples of HD cells’ coverage of the azimuthal plane, in each mouse. Rows in each matrix represent tuning curve heatmaps of individual HD cells. Tuning curve amplitudes are normalized.

**Extended Data Figure 2.**
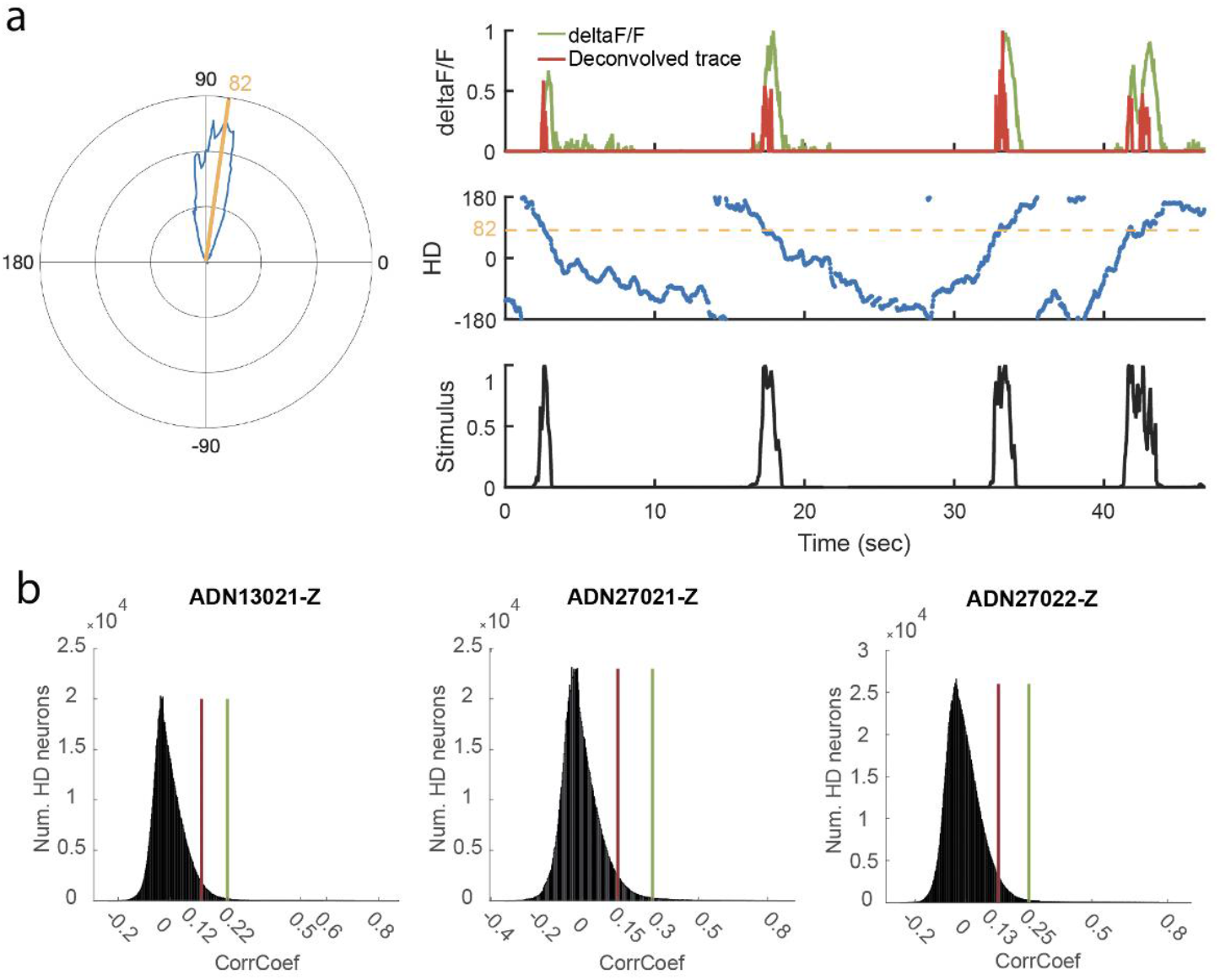
Identification of HD neurons with calcium imaging. **a**. Left: Example polar tuning curve for a HD neuron. Yellow line indicates direction of maximum firing rate (i.e. preferred firing direction (PFD)). Firing rates are occupancy normalized. Right: Top: An example calcium signal deltaF/F (green) from one HD neuron and the resulting deconvolved trace (red). Both traces were normalized. Middle: Measured head-direction. Bottom: The extracted stimulus signal of the HD neuron’s PFD. Peaks indicate instances of the animal facing the particular PFD. The deconvolved signal is cross-correlated with the stimulus signal in order to obtain the Pearson’s correlation coefficient which reflects the degree of HD tuning of the cell (r=0.85 in the case of the current example). **b**. Distributions of the Pearson’s correlation coefficients after 1000 circular-shift shuffles of the firing rate signals (smoothed deconvolved traces) of all HD neurons, in each mouse. Red and green vertical lines indicate the 95th and 99th percentiles, respectively. Data includes 10 baseline recordings of 3 minutes each for every mouse. Of all recorded cells, ~94% met the 95th percentile selection criterion while ~83% met the 99th percentile selection criterion.

**Extended Data Figure 3.**
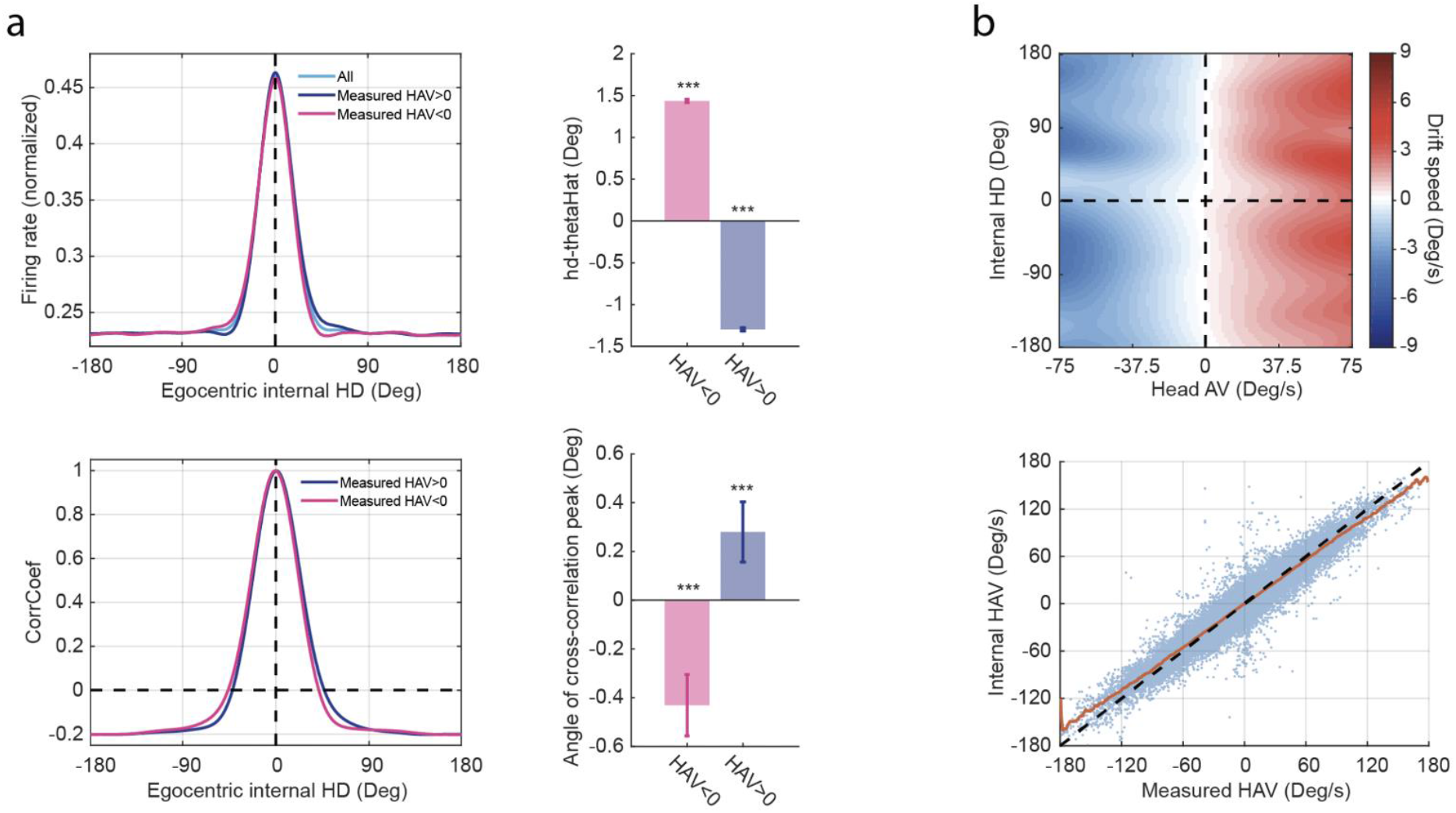
Anticipatory behaviour and drift-speed pattern during baseline. **a**. Top row: Mean bump of activity (N=42 baseline epochs) divided between positive (blue) and negative (pink) head angular velocities (HAV). Bar graph: Mean difference between measured and decoded HD (Wilcoxon signed rank test: HAV<0: p~0, Z=83.71; HAV>0: p~0, Z=−76.81). Bottom row: Mean cross-correlation (N=42 baseline epochs) of the mean bump of activity, per epoch, with the mean bump of activity for positive (blue) and negative (pink) HAVs. Bar graph: Mean peak angle of cross-correlation (Wilcoxon signed rank test: HAV<0: p~0, Z=−115.24; HAV>0: p~0, Z=113.13). Both analyses show a significant amount of anticipation of future heading by the HD network. **b**. Top: Drift-speed heatmap showing an increased latency in updating the internal representation as the HAV becomes larger. Bottom: same pattern as the above, seen here in Internal HAV-versus-Measured HAV space. Notice the deviations of the mean signal (orange) from the diagonal, at high measured HAVs. Bar graphs indicate mean and SEM.

**Extended Data Figure 4.**
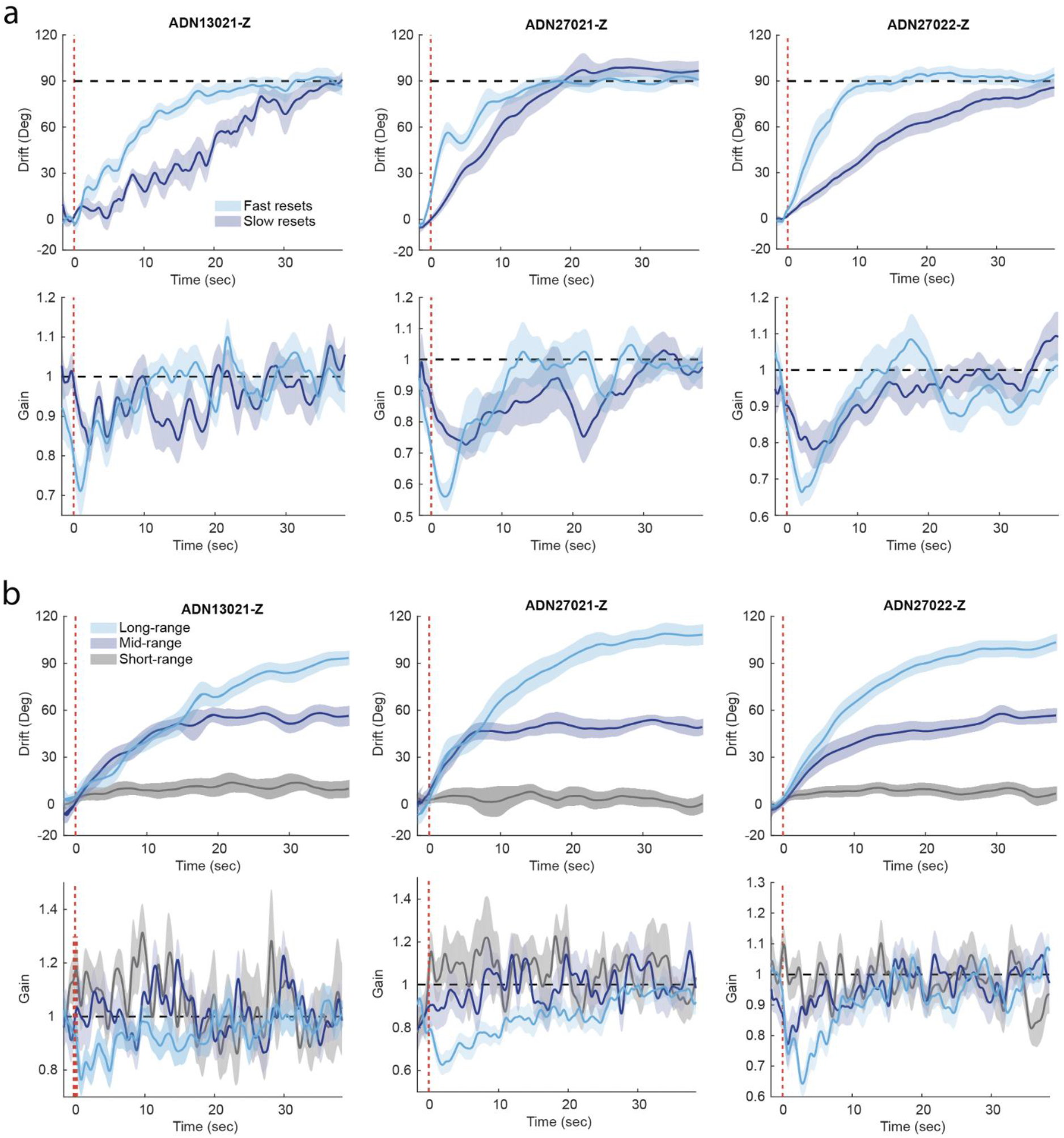
Reset behavior and gain modulation per mouse. **a**. Top row: Resets separated according to their speeds between fast (light blue) and slow (dark blue) groups, per mouse. Bottom row: Corresponding gain signals for fast and slow resets, per mouse. **b**. Top row: Resets separated according to their range between long- (light blue), mid- (dark blue) and short- (grey) groups, per mouse. Bottom row: Corresponding gain signals for long-, mid- and short-range resets, per mouse. Values are shown as Mean (solid line) and SEM (shaded area).

**Extended Data Figure 5.**
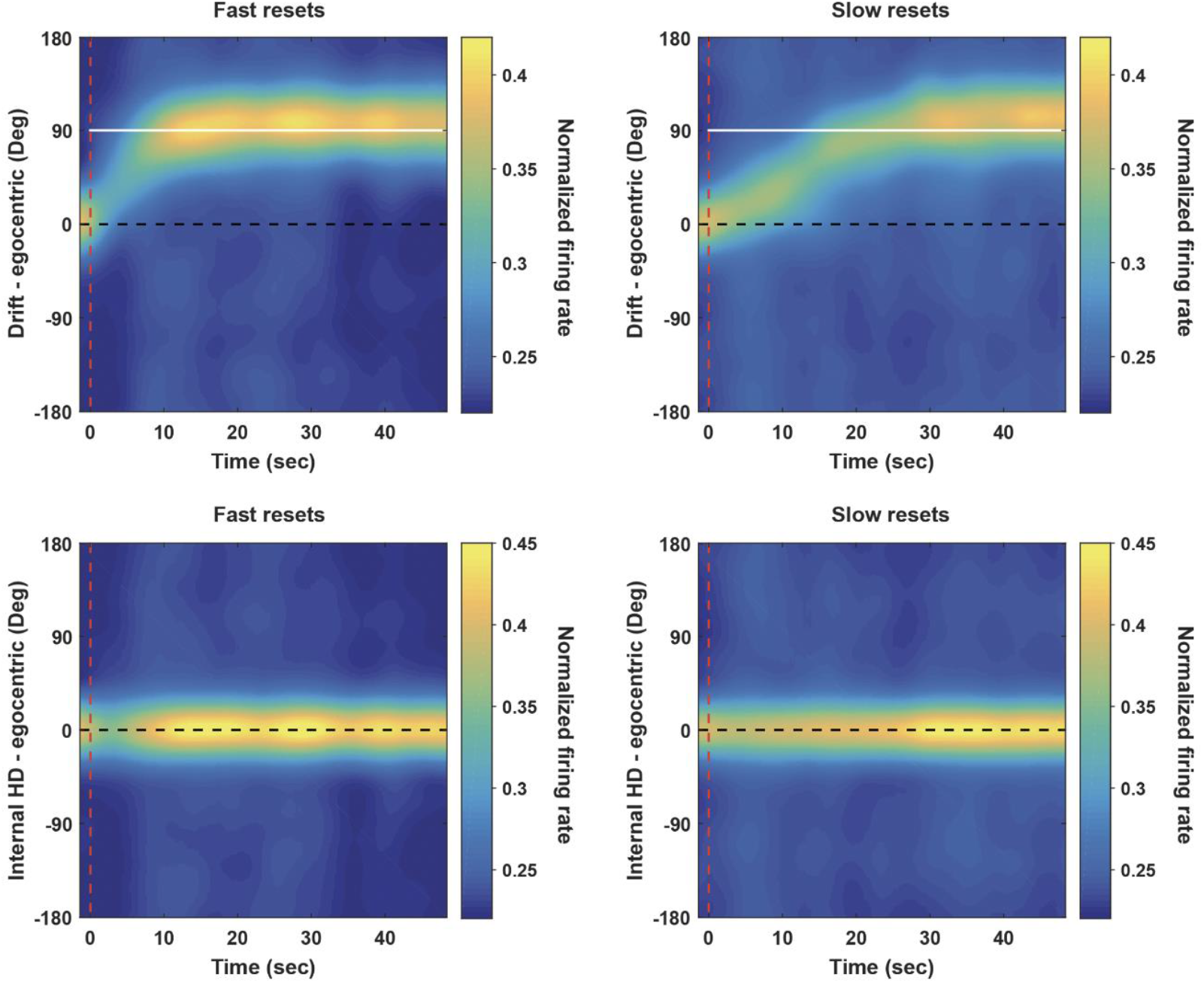
Reconstruction of the bump of activity. Averaged heatmaps of the bump of activity during fast (left column) and slow (right column) resets (same data as in Fig2 H, I). Data is presented in the egocentric reference frame, without drift adjustment (top row) and with drift adjustment (bottom row) showing, in both cases, no additional bumps outside the main activity packet. Dashed red line indicates cue-onset, while white horizontal line at 90° is for reference. Firing rates are normalized.

**Extended Data Figure 6.**
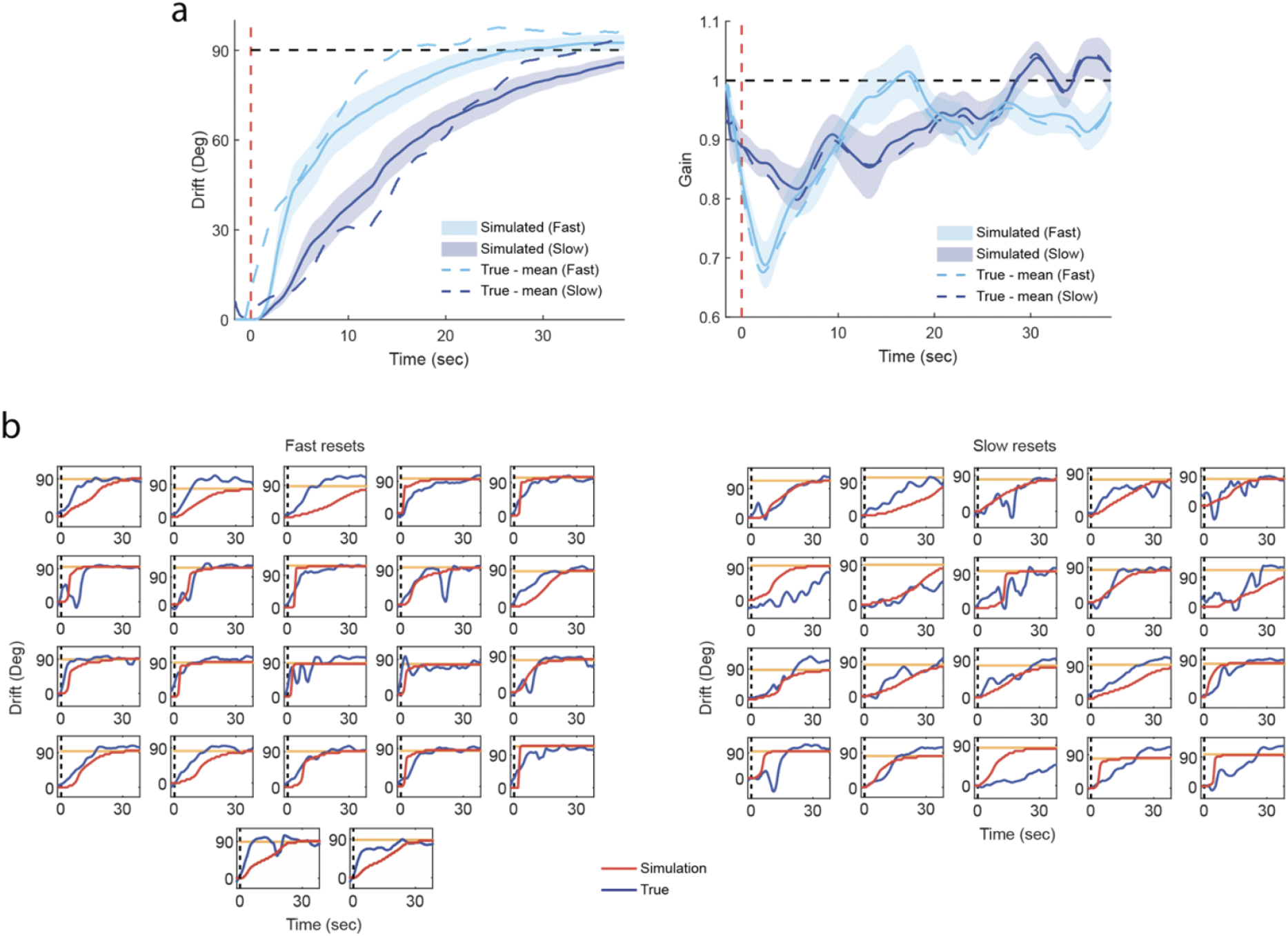
Simulation results of the HD network model and prediction of drift speed based on input gain. **a**. Left: Mean simulated reset signals for fast (light blue) and slow (dark blue) groups. Right: Mean simulated gain signals for the same groups. Values are shown as Mean (solid line) and SEM (shaded area). Dashed signals represent means of ground-truth data. **b**. Individual examples of simulation predictions (red lines) for fast and slow reset groups, plotted against actual resets (blue lines). Yellow lines indicate cue location. Amplitudes are relative to angles at cue-onset (dashed black line).

**Extended Data Figure 7.**
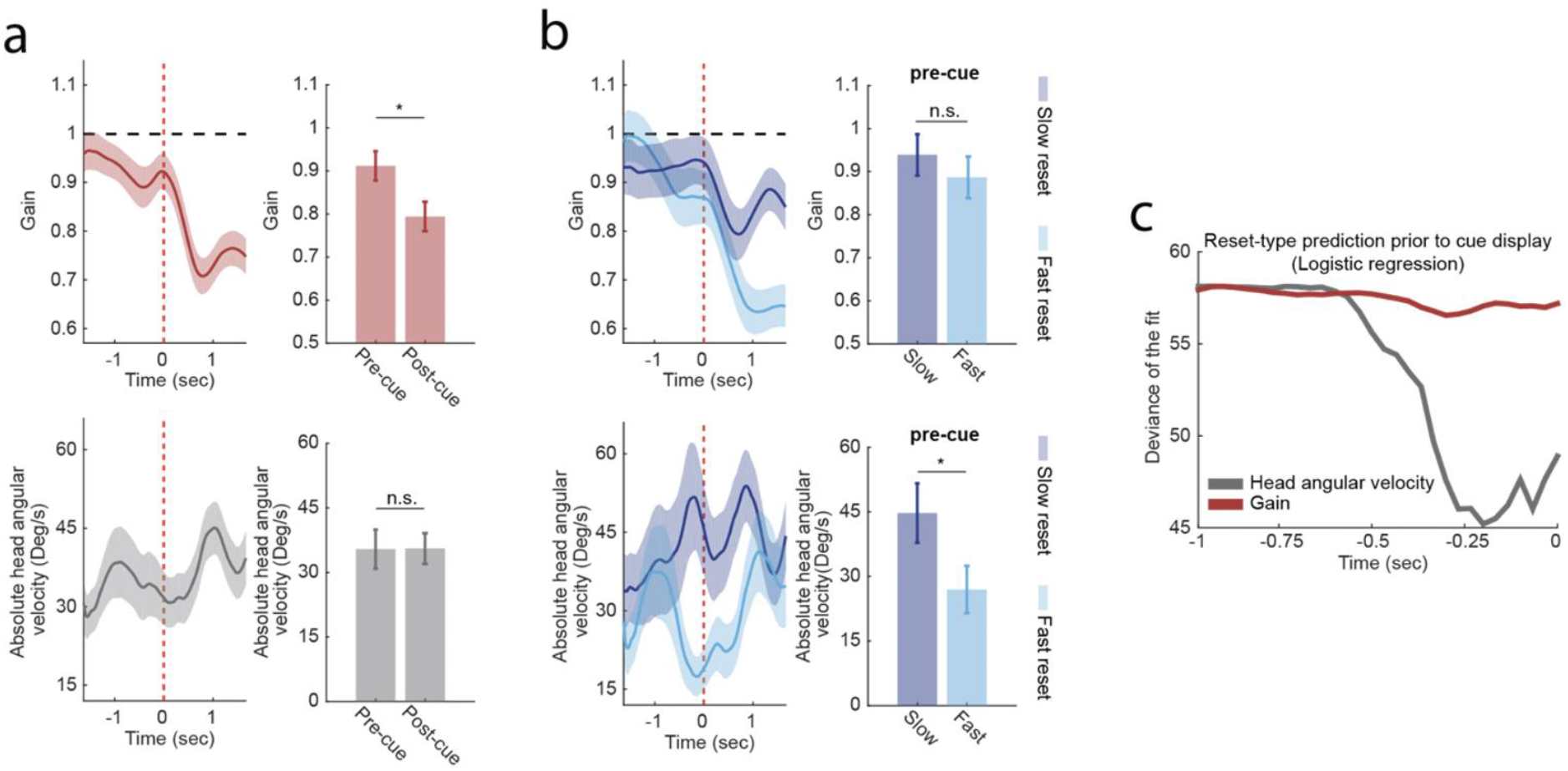
Animal behavior, prior to cue display, is predictive of reset speed. **a**. Triggered average of gain shows a sharp decrease following cue display, for 90°-centered resets ([70°:110°] range) (Wilcoxon rank sum test: average gain 1-second pre-cue versus average gain 1-second post-cue: p = 0.0228, Z = 2.28) (Top). However, overall absolute head angular velocity (aHAV) does not seem to differ before and after cue display (Wilcoxon rank sum test: average aHAV 1-second pre-cue versus average aHAV 1-second post-cue: p = 0.6259, Z = 0.49) (bottom). **b**. Separation of signals in a. between fast and slow resets shows similar gain amplitudes over a 1-second interval prior to cue display (Wilcoxon rank sum test: p = 0.3580, Z = 0.92) (Top). However, aHAV is lower for fast resets compared with slow resets, over the same period (Wilcoxon rank sum test: p = 0.0294, Z = 2.18) (Bottom). **c**. Head angular velocity becomes more predictive of reset type closer to the moment of cue-display when compared with prediction performance based on gain amplitudes within the same time interval. Deviance of the fit is used as defined in Matlab’s *mnrfit* function for logistic regression. Data shown is same as in Fig2. g. Time dependent signals, in **a** and **b**, are shown as mean (solid line) and SEM (shaded area) and bar graphs indicate mean and SEM

**Extended Data Figure 8.**
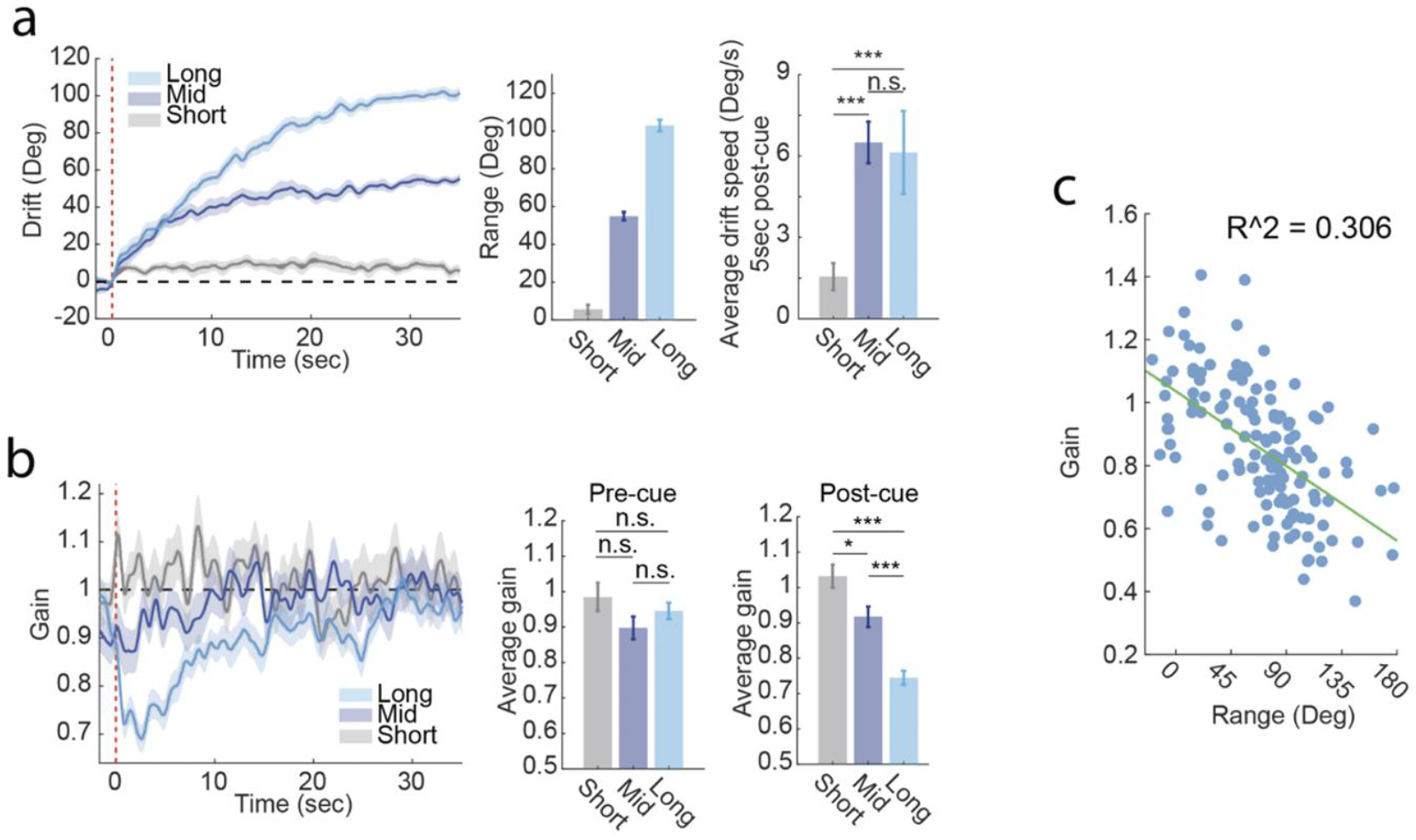
Relationship between reset range and gain modulation. **a**. Mean drifts for short- (grey; n = 27), mid- (dark blue; n = 40) and long- (light blue; 67) range reset-groups showing non-significant difference in drift-speeds between mid- and long-range groups (Wilcoxon rank sum test: Short-Mid: p=4.19e-5, Z=4.10; Short-Long: p = 7.73e-5, Z = 3.95; Mid-Long: p = 0.62, Z = 0.50; 150 frames (~5s) post-cue) **b**. Network gains for the short-, mid- and long-ranges have similar amplitudes prior to cue-display (Wilcoxon rank sum test: Short-Mid: p = 0.1174, Z = 1.57; Short-Long: p = 0.32, Z=1.00; Mid-Long: p=0.2984, Z=1.04; 50 frames (~1.67s) pre-cue), yet they exhibit gradual decrease following cue-display (Wilcoxon rank sum test: Short-Mid: p = 0.0129, Z = 2.49; Short-Long: p=2.6876e-9, Z=5.95; Mid-Long: p = 1.2130e-5, Z = 4.38; 150 frames (~5s) post-cue). **c**. Relationship between average gain and reset range. Each dot represents a correct reset (n = 134). The R-squared value corresponds to a linear regression model fit (green line). All CW sessions have been reflected across the x-axis and transformed into CCW ones. Time-dependent signals are shown as mean (solid line) and SEM (shaded area) and bar graphs indicate mean and SEM.

**Extended Data Figure 9:**
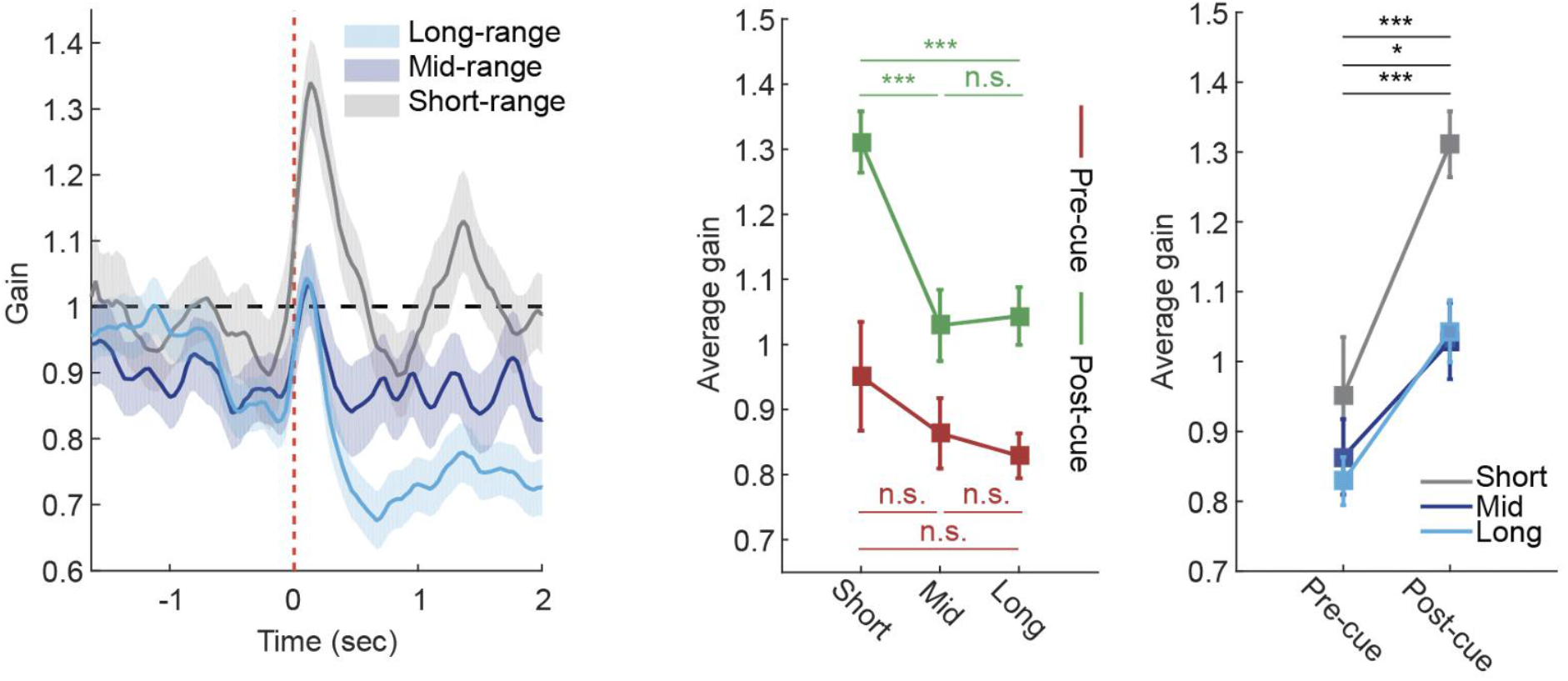
Rapid gain spikes can be seen shortly after cue-display, in the three reset-range groups (same data as in Fig. 2m shown at higher temporal resolution). All reset ranges start at similar amplitudes at the end of the darkness period (Wilcoxon rank sum test: short-mid: p=0.3940, Z=0.85; short-long: p=0.2090, Z=1.26; mid-long: p=0.4686, Z=0.72). Following cue-display, each group exhibits a brief gain increase (5 frames (~150ms) pre-cue vs 5 frames (~150ms) post-cue: Wilcoxon rank sum test: short: p=6.9690e-4, Z=3.39; mid: p=0.0369, Z=2.09; long: p=2.6898e-4, Z=3.64). These gain spikes are largest for the short-range group (Wilcoxon rank sum test: short-mid: p=4.4888e-4, Z=3.51; short-long: p=1.8600e-4, Z=3.74; mid-long: p=0.9326, Z=0.08). Time-dependent signals are shown as mean (solid line) and SEM (shaded area) and error bars indicate mean and SEM.

**Extended Data Figure 10.**
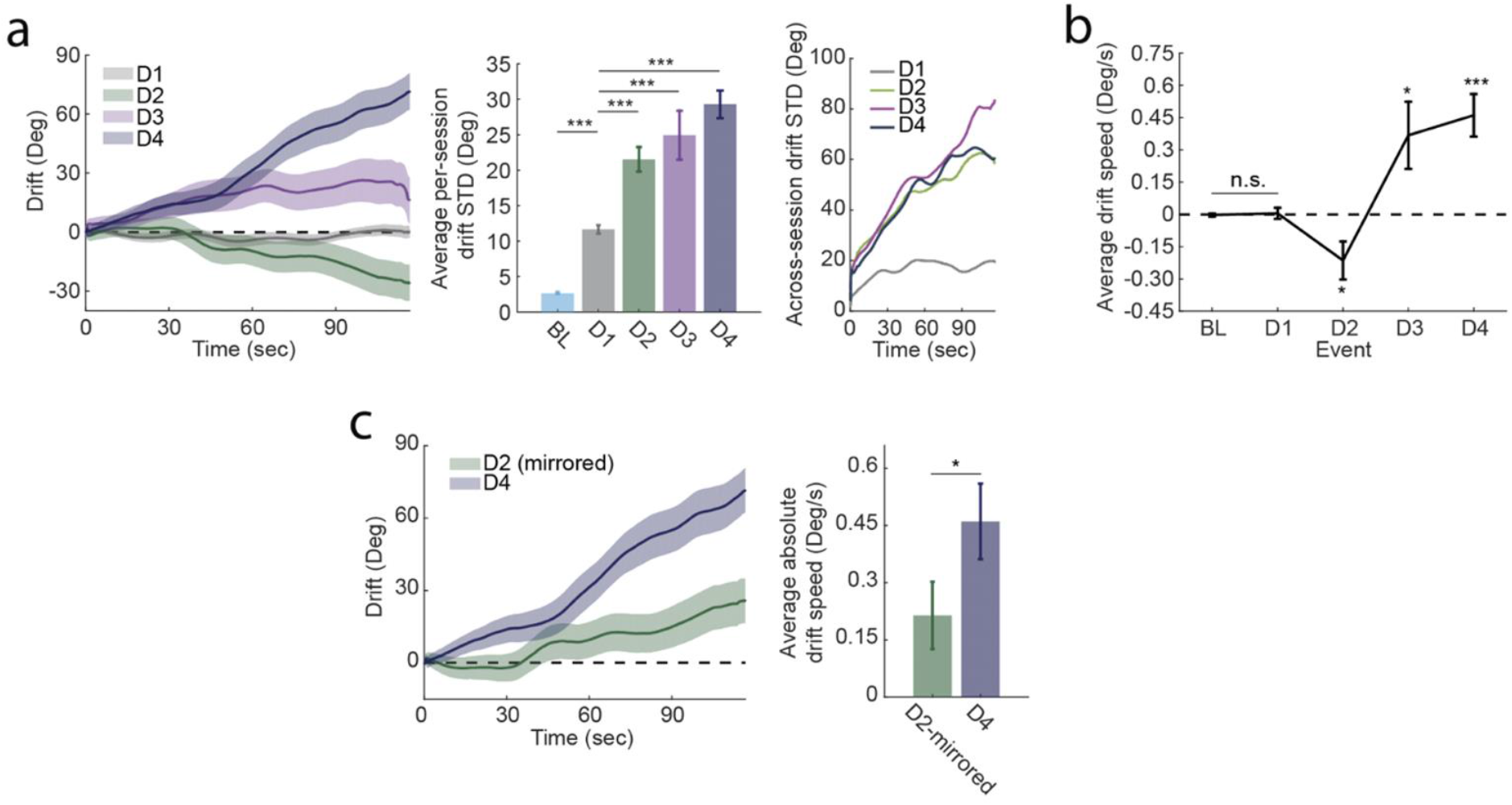
Distinct drift patterns across darkness periods. **a**. Drift variability increases significantly following a reset (D2, D3 and D4) in comparison with D1 (Mean drift STD compared across darkness epochs: Wilcoxon rank sum test: BL-D1: p = 3.1214e-15, Z = 7.89; D1-D2: p = 1.1477e-6, Z = 4.86; D1-D3: p = 8.3761e-5, Z = 3.93; D1-D4: p = 5.6600e-11, Z = 6.55). Drift STD also increases with time after a reset (D2, D3 and D4) while it remains constant following baseline (D1). **b**. Mean drift-speed in each darkness epoch. Systematic biases depend on prior cue-event. (Wilcoxon rank sum test: BL-D1: p=0.1250, Z=1.53; Wilcoxon signed rank test: D2: p = 0.0168, Z = −2.39; D3: p = 0.0313, Z = 2.15; D4: p = 2.9929e-4, Z = 3.62). **c**. Comparison between drifts in D2 and D4 of the 90°-cue-shift experiment. Although the two events are experimentally symmetric to each other w.r.t baseline, drifts in D4 appear to have larger biases (in absolute value terms) than D2. Left: Mean drift signals, in D2 (dark blue) and D4 (light blue). Drifts in D2 have been mirrored across the 0°-line for comparison purposes. Values are shown as Mean (solid line) and SEM (shaded area). Right: Comparison between average drift speeds, in D2-mirrored (Light blue) and D4 (dark blue) (Wilcoxon rank sum test: p=0.0184, Z=2.36). Time-dependent signals are shown as mean (solid line) and SEM (shaded area). Bar graphs, in a and c, and error bars, in b, indicate mean and SEM.

**Extended Data Figure 11.**
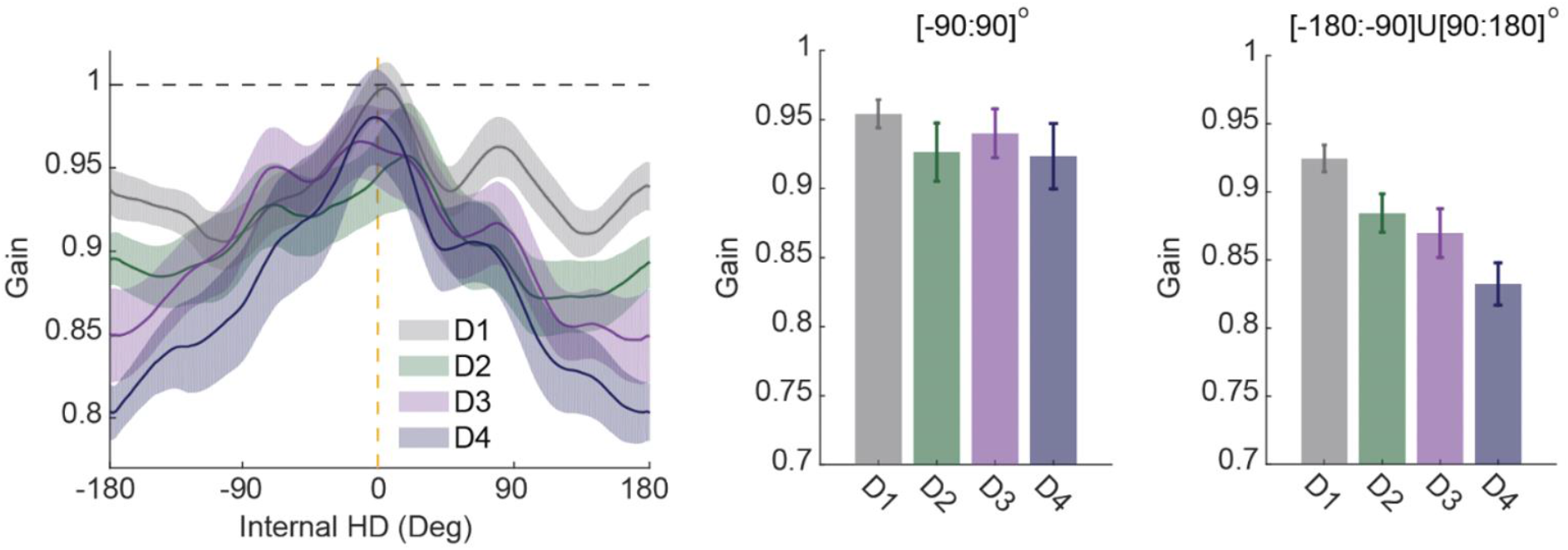
Time dependent changes in gain tuning curve. Average gain tuning curves across darkness periods showing gradual decrease of network gain away from the internal cue location (dashed yellow line) from D1 to D4. Tuning curves are shown as mean (solid line) and SEM (shaded area) and bar graphs indicate mean and SEM.

**Extended Data Figure 12.**
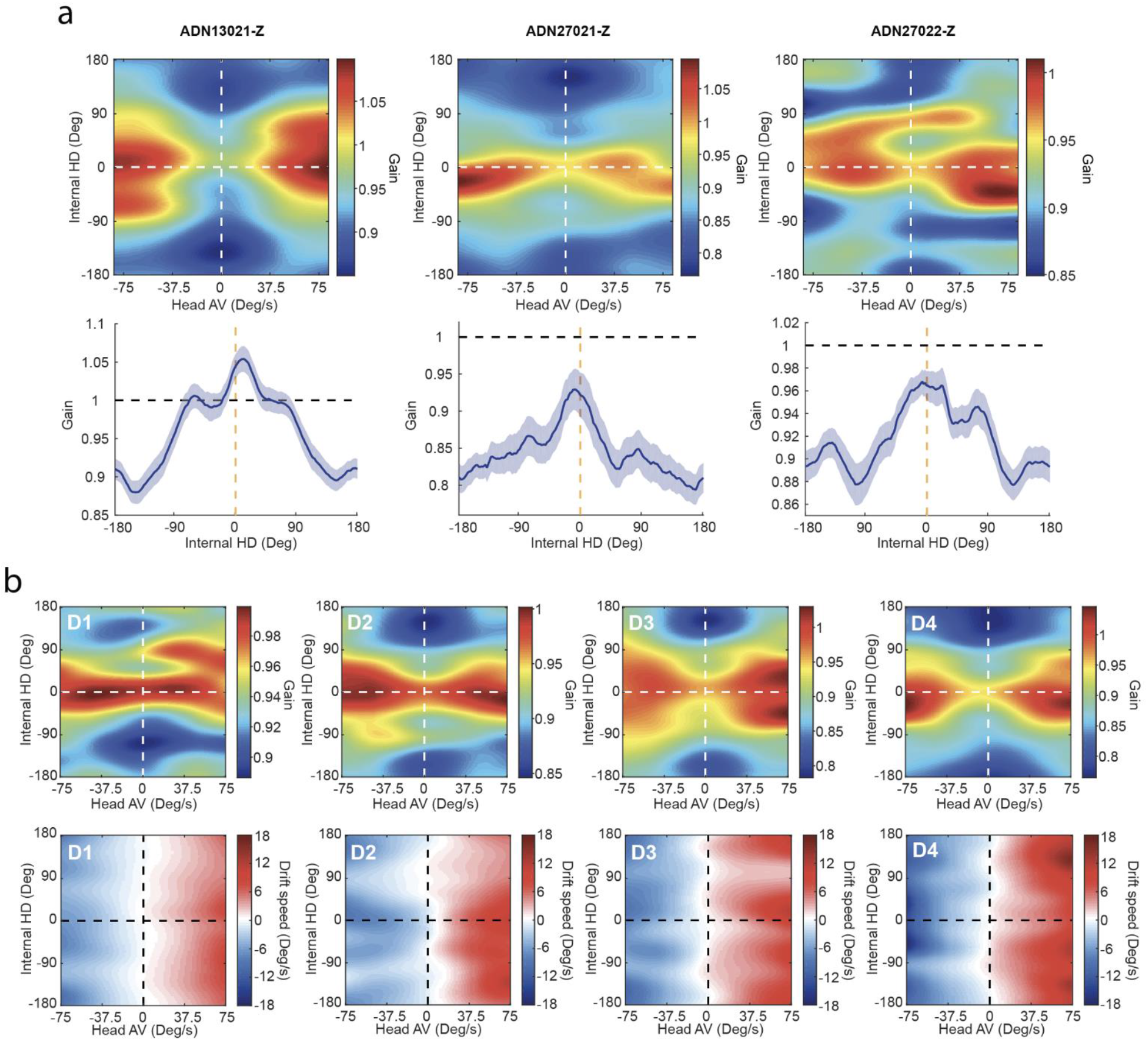
Network gain patterns across mice and darkness epochs. **a**. Network gain during darkness shown as heatmaps (top row) and tuning curves (bottom row), per mouse. In both cases, data is averaged across sessions and darkness epochs (D1 to D4) of the 90°-cue-shift experiment. Values for the tuning curves are shown as Mean (solid line) and SEM (shaded area). **b**. Top row: Network gain heatmaps showing same data as in (A) split (from left to right, respectively) across the different darkness epochs D1 to D4 of the 90°-cue-shift experiment. Bottom row: Drift speed heatmaps showing a consistent pattern, yet with varying amplitudes, across darkness epochs D1 to D4. No obvious effect of the gain landscape can be seen in these patterns.

**Extended Data Figure 13.**
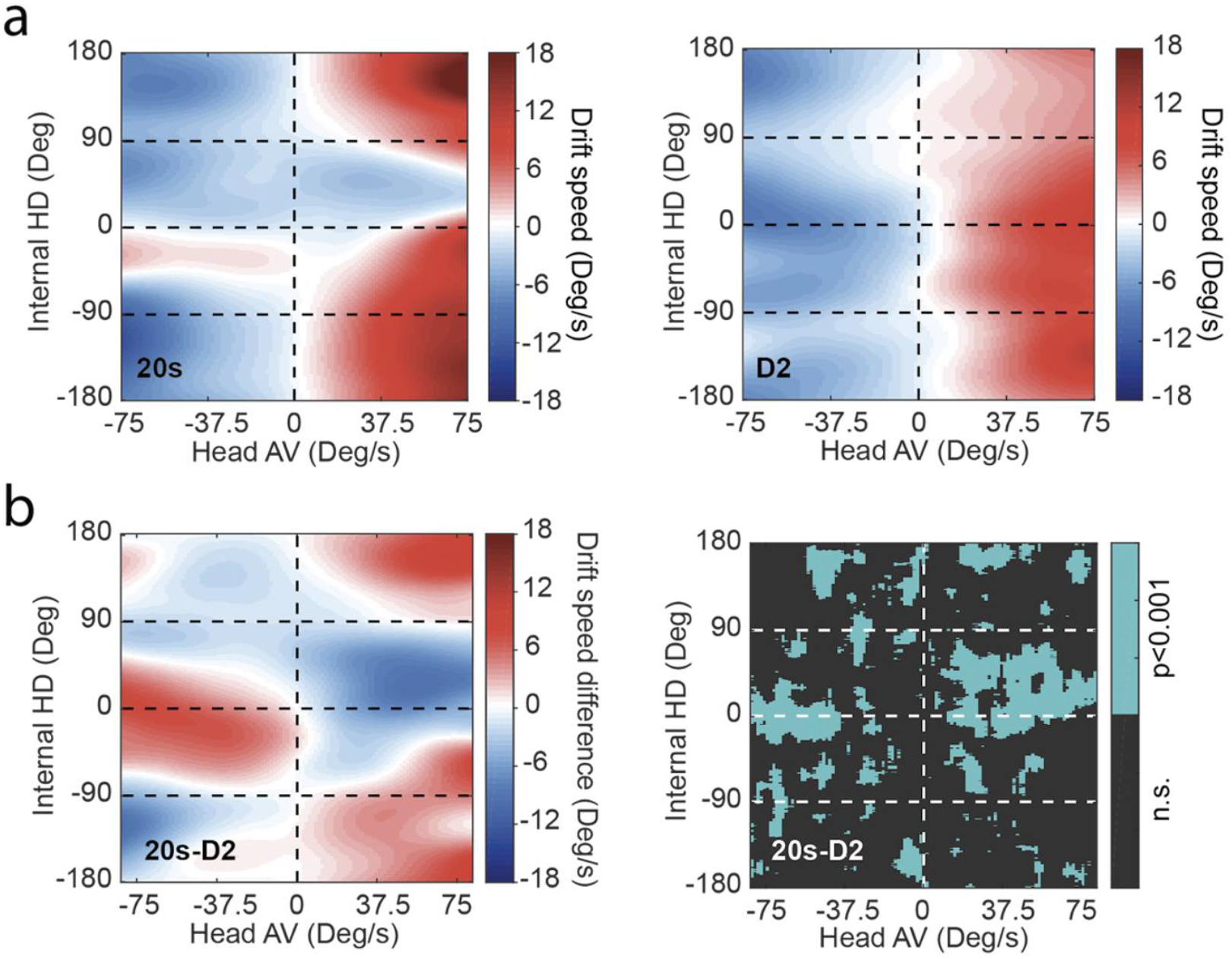
Distortion of the drift speed landscape during reversion. **a**. Drift-speed heatmaps. Left: 20s-cue-exposure experiment (n = 43). Right: D2 of the 2m-cue-exposure experiment (n = 35). **b**. Left: Drift-speed difference (same data as in **e**) showing a significant distortion of the pattern seen in the first experiment around the internal location of the cue. Right: p-value matrix for data in left (Wilcoxon rank sum test; pixels where p>0.001 were marked as NaN).

**Extended Data Figure 14.**
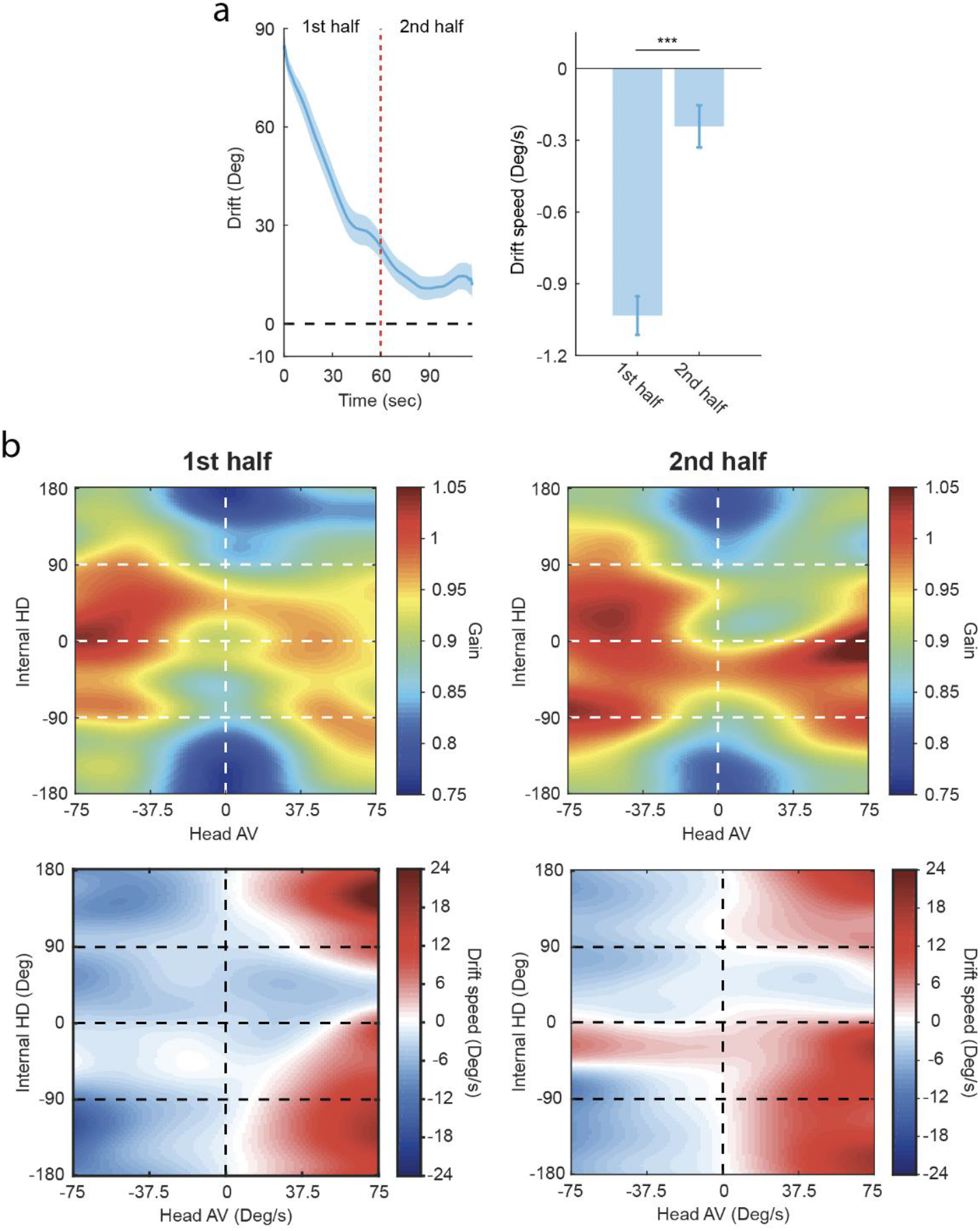
The two stages of reversion in the 20s cue-exposure experiment. **a**. Mean drift signal during reversion (N=43). Dashed yellow line divides the darkness period in two halves with contrasting states of the HD network: drifting (1st half) and stabilizing (2nd half). Data shown as mean (solid line) and SEM (shaded area). Bar graph: Comparison of mean drift-speeds between the first and second halves of the darkness period (Wilcoxon rank sum test: p=3,5802e-8, Z=5.51). Data shown as mean and SEM. **b**. Top row: Heatmaps of network gain during the first (left) and second (right) halves. Bottom row: Heatmaps of drift-speed during the first (left) and second (right) halves showing state-dependent distortions of the drift-speed pattern.

**Extended Data Figure 15.**
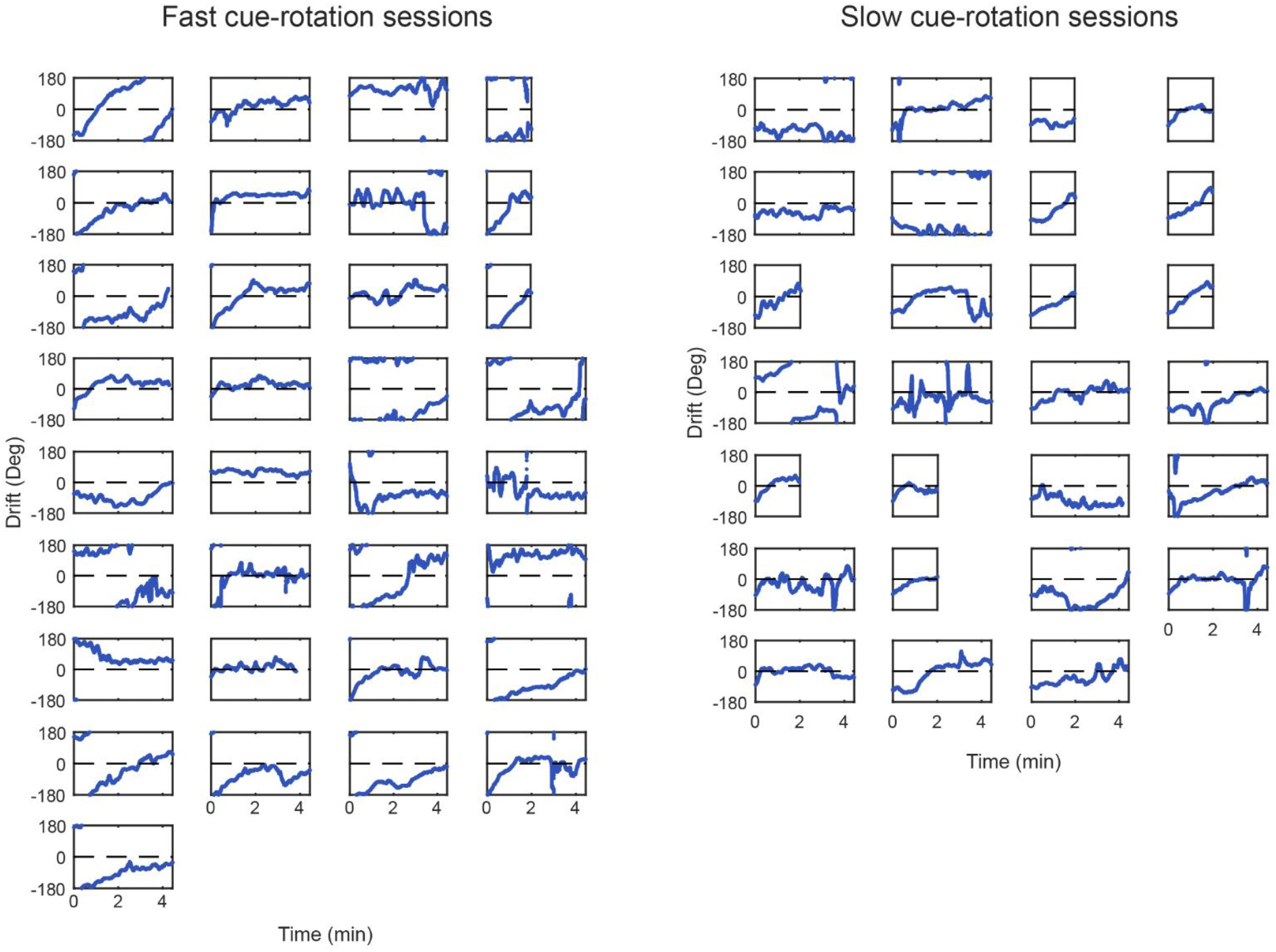
Individual examples of drift biases during darkness following a continuous fast (left group) or slow (right group) cue-rotation. At the beginning of each cue-rotation epoch, the visual cue was displayed at the same location as that of baseline. It, then, keeps rotating for seven minutes. At the end of the seven minutes and depending on the speed of rotation, the cue would either reach ±180° or −90°. The drift signal is, thus, expected to start within a close range of these two directions, during the second darkness epoch. However, in some cases, drifts during the first darkness epoch were large enough so that the initial anchoring to the rotating cue occurred considerably far from baseline. This caused the drift signal, during the second darkness, to start further away from the expected location. For our comparison, in Fig. 5d, we limited our analysis to drifts starting at [−180:−145]U[145:180]° for the fast cue-rotation group and [−125:−55]° for the slow cue-rotation group, in order to study the effects across sessions with similar stability during baseline (total N=44 out of 60).

**Extended Data Figure 16.**
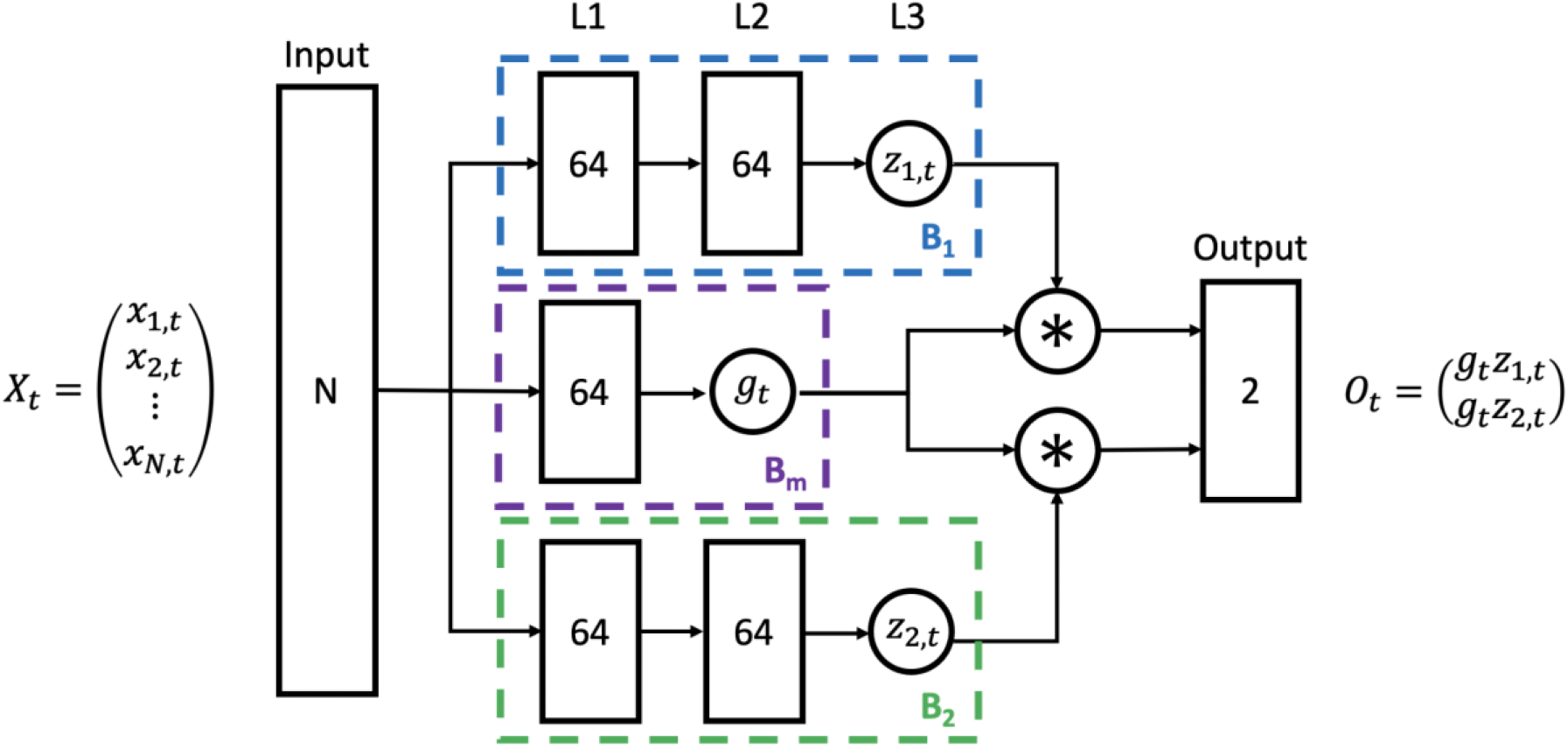
Diagram of the artificial neural network used to project high dimensional neural activity onto 2D polar space. Numbers inside each box correspond to the unit count. All activation functions are ‘relu’ except for nodes *z_1,t_* and *z_2,t_* where the activation function is ‘tanh’. In all layers, we apply *L*_2_ regularization with regularization factor 0.001. Input data, from *N*ADN neurons, are normalized.

## Supplementary material: Attractor network model

The goal of our model is to propose a potential neural mechanism of drift control, in the HD system, through gain modulation. The neuronal model is based on that described in Redish et al. (1999)^45,54^. However, we made important changes to the network design which will be detailed below.

### Model inputs

*α_in_*: Time-series of true network gain

*AV*: Time-series of true head angular velocity (AV), in degrees/frame

*θ_cue_*: Angle of current cue on display, in degrees

*θ_HD_*: Animal’s true head-direction, in degrees

*D_ini_*: Initial drift offset, in degrees

*L*: Binary time-series indicating cue display (0:cue-off, 1:cue-on)

### Model outputs

*α_sim_*: Time series of simulated network gain of the HD-neuron layer

*D_sim_*: Time series of simulated drift signal, in degrees

## 1 Network design

Our attractor network is composed of three layers (pools): (1) The HD layer, (2) the inhibition layer and, (3) the conjunctive AV-by-HD layer which itself can be divided into two sub-layers: (3a) the CW-AV-by-HD layer and (3b) the CCW-AV-by-HD layer. Extra-network input comes in the form of a visual layer, AV cells (CW and CCW) of the vestibular system as well as a global modulation source (i.e., gain cell).

### 1-1 Generation circuit

First, we show how the generation circuit (layers (1), (2) and (3)) can produce a stable HD representation. Model figures 1, 2 and 3 show the connections between the layers. Our model can maintain a unique bump of activity (at the HD layer) through constant input from the vestibular system (AV-by-HD layer) combined with lateral inhibition from the inhibitory layer. Our choice of departing from standard HD-network models that use recurrent excitation at the HD layer to generate a unique bump of activity is motivated by anatomical and physiological studies of the HD circuitry, in rodents. We know that two types of AV cells exist in the vestibular system: symmetric and asymmetric AV cells (CW-AV cells and CCW-AV cells)^14,55^. While the symmetric AV cells increase their firing rate proportional to the head AV, regardless of direction, the asymmetric AV cells’ activity increases only in one direction and decreases in the other. Interestingly, both asymmetric AV cells’ subtypes (CW and CCW) appear to fire at higher rates than minimum values, albeit with equal amounts, when the animal’s head is not moving. We also know the involvement of inhibition in the generation of HD-cell activity from previous studies that showed the connections in the downstream pathway of the generation circuit (DTN→LMN) are largely GABAergic^56,57^. Additionally, there is no anatomical evidence for the existence of recurrent excitation neither in the generation circuit (DTN and LMN) nor in the thalamus (ADN). Therefore, achieving a stable representation by combining the activity from the vestibular system together with a lateral inhibition (as in the current model) appears more biologically plausible.

#### Neuronal dynamics

We use a similar approach to Redish (1996) to model the firing activity of every neuron in the HD, inhibitory and AV-by-HD layers^54^. Each postsynaptic unit’s response is governed by three equations:

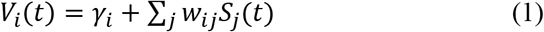

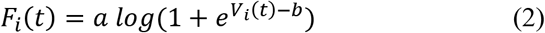

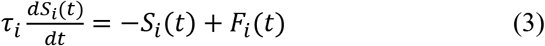

Where, *V_i_* is the voltage of postsynaptic neuron *i*, *γ_i_* is the tonic inhibition term, *ω_i,j_* is the synaptic weight between postsynaptic neuron *i* and presynaptic neuron *j*, *S_j_* is the synaptic drive of presynaptic neuron *j*, *F_i_* is the activation function of postsynaptic neuron *i*, *dt* is the time step and, *τ_i_* is the time constant defining the decay rate of postsynaptic potential (PSP). Equations (1) and (3) follow directly from Reddish (1996)^54^. However, we opted for a more biologically plausible activation function *F_i_*, in (2), following Zhang (1996)^58^ which also has the advantage over the hyperbolic tangent (used in Redish (1996)) of preventing saturation issues at high activity levels. Both *a* and *b*, in (2), are optimization parameters that control the scale and shift of the activation function, respectively.

Our model is composed of the same number of units (neurons), *N*, per layer/sublayer. To understand how this works, it is useful to divide the horizontal plane into *N* equally distant directions which we will refer to as preferred firing directions (PFDs). We, then, assign to every unit, in each layer/sublayer, a unique PFD. We refer to neurons with the same PFD but belonging to different layers as ‘counterparts’.

#### Interactions between the HD and the inhibitory layers

Each inhibitory unit sends projections to all HD neurons. To ensure maintenance of the characteristic bell shape of the activity packet (at the HD layer), the synaptic weights are determined by a Gaussian kernel such that they become stronger with the increase in PFD distance between the inhibitory neuron and its target HD neuron. Effectively, when active, an inhibitory neuron causes minimal decrease in firing activity within a close neighborhood of its counterpart HD neuron while it engenders maximal inhibition on distant HD units.

The connection weight of the projection from inhibitory neuron *i* onto HD neuron *j* is, thus, given by:

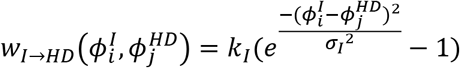

Where, 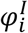 and 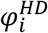 are PFDs of inhibitory neuron *i* and HD neuron *j*, respectively, *k_I_* is a scale factor and, *σ_I_* is the standard deviation of the weight distribution.

On the other hand, each inhibitory neuron receives an excitatory back projection from its unique counterpart HD neuron. This ensures only a subset of the inhibitory pool (counterpart of the activity packet) is active, at any given time, resulting in lateral inhibition of HD neurons outside the activity packet

**Model Figure 1:**
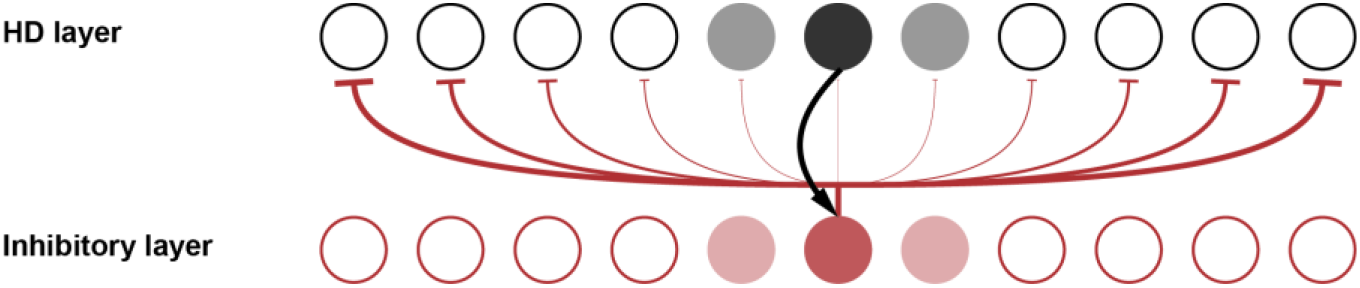
Connections between the HD layer and the inhibitory layer. Triangular arrowhead indicates excitatory projection. Flat arrowhead indicates inhibitory projection. Color gradients indicate the level of activity for each neuron (i.e., opacity increases with firing activity). Arrow thickness indicates synaptic strength (i.e., thickness increases with synaptic weight). For clarity, we only show projections from the unit with highest activity, at each layer

#### Interactions between the HD and the AV-by-HD layers

Like the inhibitory layer, every AV-by-HD neuron sends direct projections to all HD neurons. However, in this case, the synaptic weights are determined by a Gaussian kernel that peaks at an offset location w.r.t the counterpart HD neuron. Concretely, an AV-by-HD unit provides the highest excitation either rightwards (for a CW-AV-by-HD neuron) or leftwards (for a CCW-AV-by-HD neuron) of the counterpart HD neuron. This configuration has the advantage of allowing more flexibility and fine tuning in the calibration of the system (i.e., matching vestibular input to visual flow). The amount of offset is kept the same (in absolute value terms) between CW and CCW AV-by-HD units, which ensures balanced input from both sublayers when the animal’s head is not rotating.

The connection weight of the projection from AV-by-HD neuron *i* onto HD neuron *j* is given by:

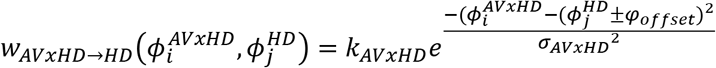

Where, 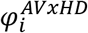 and 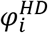 are PFDs of AV-by-HD neuron *i* and HD neuron *j*, respectively, *k_AVxHD_* is a scale factor, *σ_AVxHD_* is the standard deviation of the weight distribution and, *ϕ_offset_* is the angular offset between AV-by-HD and HD layers.

On the other hand, each AV-by-HD neuron receives an excitatory back projection from its unique counterpart HD neuron. This ensures only a subset of the AV-by-HD pool (counterpart of the activity packet) has higher activity rates than the rest, at any given time.

**Model Figure 2.**
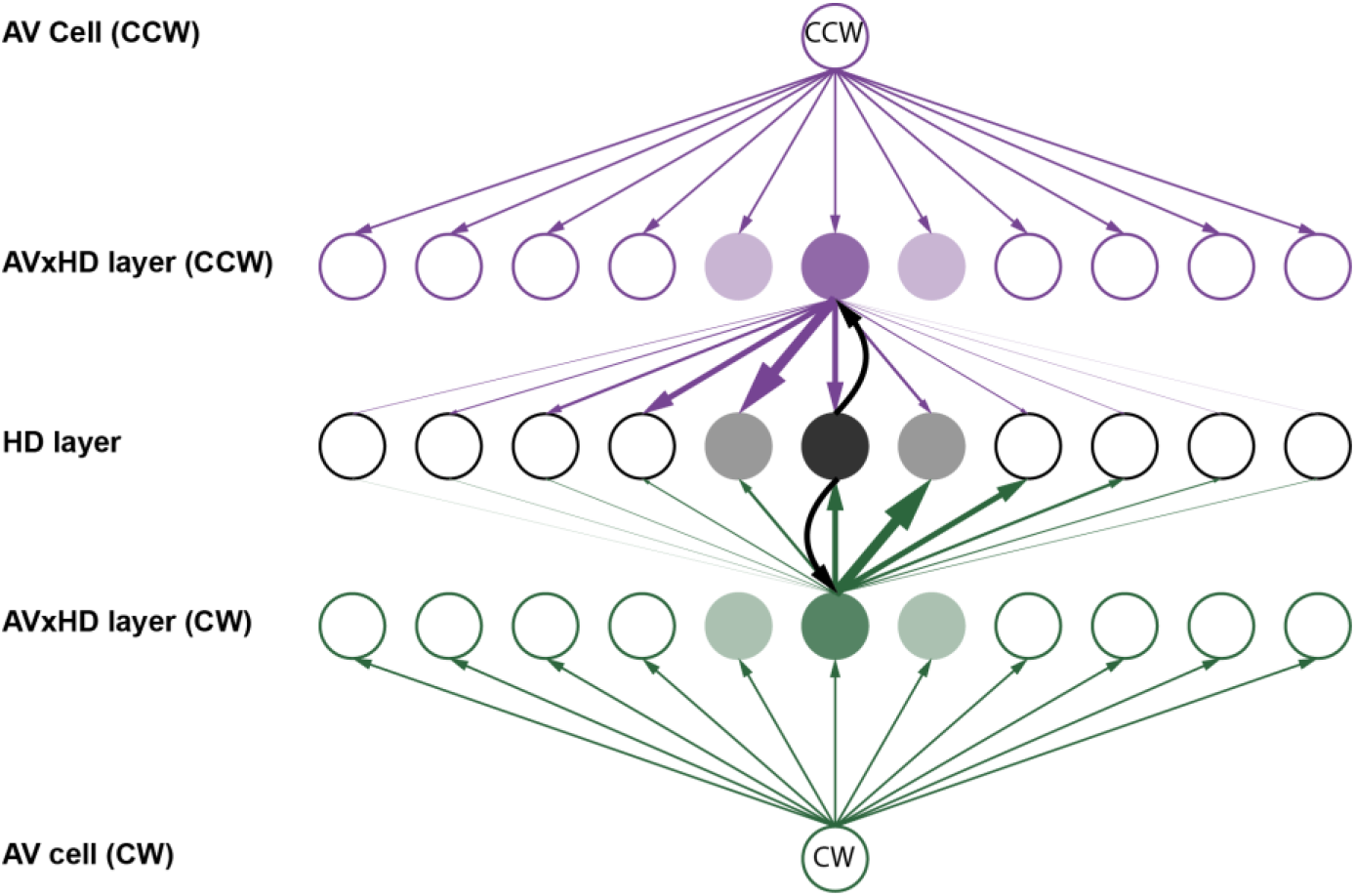
Connections between the HD layer and the AV-by-HD layers. Triangular arrowhead indicates excitatory projection. Color gradients indicate the level of activity for each neuron (i.e., opacity increases with firing activity). Arrow thickness indicates synaptic strength (i.e., thickness increases with synaptic weight). For clarity, we only show projections from the unit with highest activity, at each layer

#### Voltage calculation

With the above information considered, we can write the voltage equations for every neuron, in each layer, as follow:

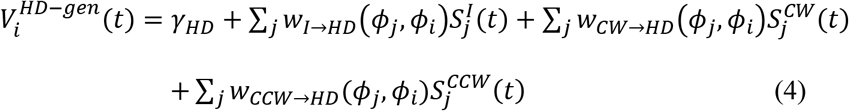

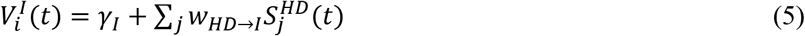

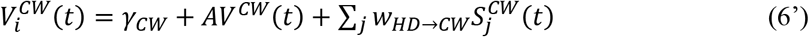

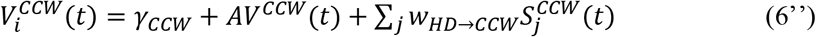

Where, 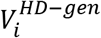 is a unit’s voltage in the HD layer, considering the generation circuit only, 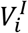 is a unit’s voltage in the inhibitory layer and, 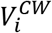 and 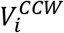 are units’ voltages in CW-AV-by-HD and CCW-AV-by-HD layers, respectively. *AV*^*CW*^ and *AV*^*CCW*^ are excitatory inputs reflecting the activity of asymmetric AV cells such that, at any given time:

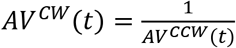

When the animal’s head is not rotating, the HD network receives equal excitation from both AV cell types such that *AV*^*CW*^(*t*) = *AV*^*CCW*^(*t*) = 1

### 1-2 Visual control of the HD network via gain modulation

#### Visual input

The visual input is provided by an extra network layer of *N* neurons. Each one of them provides an excitatory input to its counterpart HD neuron. When the visual input is available, activity on the visual layer is defined by a Gaussian kernel such that:

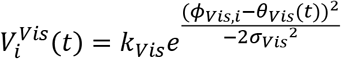

Where, *ϕ_Vis,i_* is the preferred firing direction of visual neuron *i*, *σ_Vis_* is the standard deviation of the visual Gaussian kernel and, *θ_Vis_*(*t*) is the animal’s head-direction, at time *t*, w.r.t the visual reference frame, such that: *θ_Vis_*(*t*) = *θ_HD_*(*t*) − *θ_cue_*(*t*). The input is multiplied by a constant *k_Vis_*.

#### Gain modulation

We model gain modulation as an extra-network input *g* that affects uniformly all HD neurons, which takes the form of an affine function of the experimental gain *α_in_*(*t*), such that:

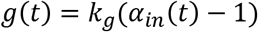

Where, *k_g_* is a positive constant. This input can be either excitatory (*α_in_*(*t*) > 1) or inhibitory (*α_in_*(*t*) < 1).

#### Integration of visual input and gain modulation

Both the visual and the gain modulation inputs constitute the downstream input to the HD layer. The final form of the voltage at a given HD neuron thus becomes:

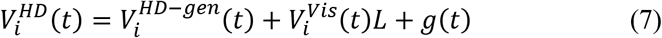

Where, *L* indicates whether the visual cue is available. It has been demonstrated, in Jackson & Redish (2003), that adding an extra-network excitatory input (equivalent to the visual input, in our case) causes the bump of activity to shift towards the direction of highest external excitation and realigns the current HD representation with the location of the external input, regardless of any movement of the animal’s head. Depending on the strength of this input as well as its distance to the current representation, the bump can either rotate continuously and span all intermediate directions (i.e., when the external input is near the current representation) or jump abruptly to the new location (i.e., when the external input is far away from the current representation). Our model allows rotations at different speeds to occur regardless of the visual input’s strength or distance to the current representation, which better reflects our experimental data. This can be achieved through gain modulation. For example, when the *g* is negative (i.e., inhibitory), the activity on the HD layer decreases which results in a weaker lateral inhibition. This makes it easier for an external input to activate HD neurons outside the activity packet and so, cause a shift in representation (note that this is a winner-takes-all situation which means, at any given time, there can only be a unique bump of activity). Conversely, if *g* is positive (i.e., excitatory), the increased activity on the HD layer renders the activation of HD neurons outside the activity packet harder and would need a stronger external input than in the previous case (i.e., *g* < 0) to achieve a similar shift in representation.

**Model Figure 3.**
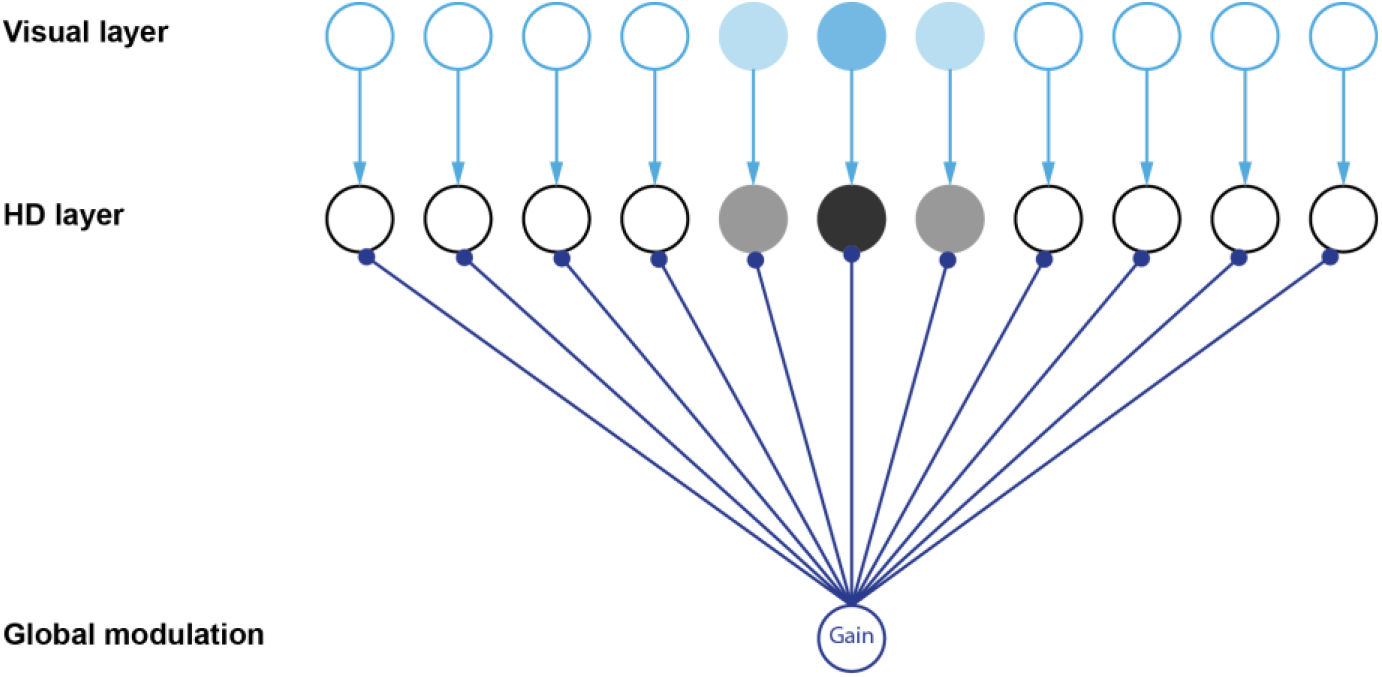
Visual control and gain modulation of the HD layer. Triangular arrowhead indicates excitatory projection. Round arrowhead indicates projection that can be excitatory (i.e., *g* > 0) as well as inhibitory (i.e., *g* < 0). Color gradients indicate the level of activity for each neuron (i.e., opacity increases with firing activity). Arrow thickness indicates synaptic strength (in this case, all synapses have similar weights).

## 2 Simulation of drift and output gain

Once all variables have been defined, we can start simulating different reset scenarios by displaying the cue at different positions while varying the input gain using experimental data. For simplicity, we assume a noise-free model. To estimate the internal HD representation, we identify the peak location of the synaptic drive on the HD layer such that:

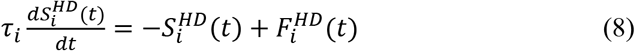

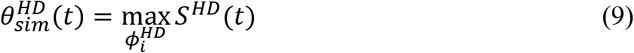

Where, *S*^*HD*^(*t*) is a Nx1 vector. To calculate the drift, *D_sim_*, we simulate two networks, in parallel, starting with the same initial conditions. The first network assumes a perfect integration of the vestibular input and does not include any visual interference or gain modulation (*g* = 0). This constitutes the reference HD, 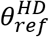 (equivalent to the measured HD, in our experimental data). The second network includes the visual input as well as the gain modulation as described, in paragraph 2-2, and from which we obtain 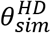. The simulated drift is simply the angular difference between the two quantities:

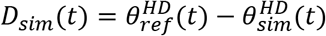

To calculate the output gain, *α_sim_*, we use the egocentric version of *S*^*HD*^, defined as:

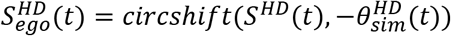

Where, *circshift* is the circular shift operator. This has the effect of bringing the activity packet to the center of the internal HD space (see example, in main text, Fig. 2E). Then, we perform a linear regression such that:

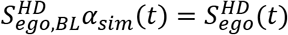

Where 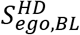 is a Nx1 vector corresponding to the egocentric synaptic drive profile of the HD layer, in baseline simulation.

## 3 Optimization

The goal of the optimization is to determine the parameter values that minimize the classification error of the drift signal (i.e., fast vs slow) while ensuring the output gain does not deviate from the input gain. Note that the output gain is not guaranteed to be the same as the input gain, due to the complex interactions between the multiple layers and the intrinsic firing properties of each unit, in the network.

We define the distance between output drift and the mean input drift, for both fast and slow resets, as follows:

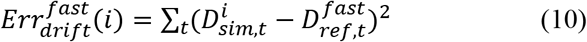

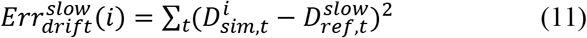

Where, 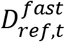 is the mean drift signal for fast-reset examples of the training set and 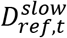 is the mean drift signal for slow-reset examples of the training set.

We then determine the simulated reset type of example *i* such that:

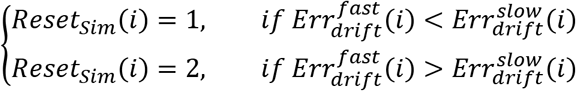

Similarly, we define the distance between output and input gains:

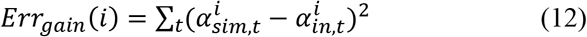

Finally, we use a search algorithm to find the optimal parameters such that:

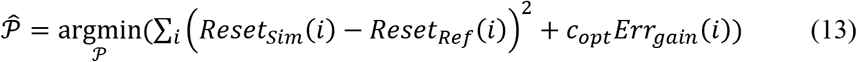

Where, *P* is the set of parameters to be optimized, *Reset_Ref_* is a 1-D vector indicating the type of the training reset examples (i.e., 1=fast and 2=slow) and *c_opt_* is a penalizing term that determines how much weight is attributed to the minimization of input-output gain error.

**Model Figure 4.**
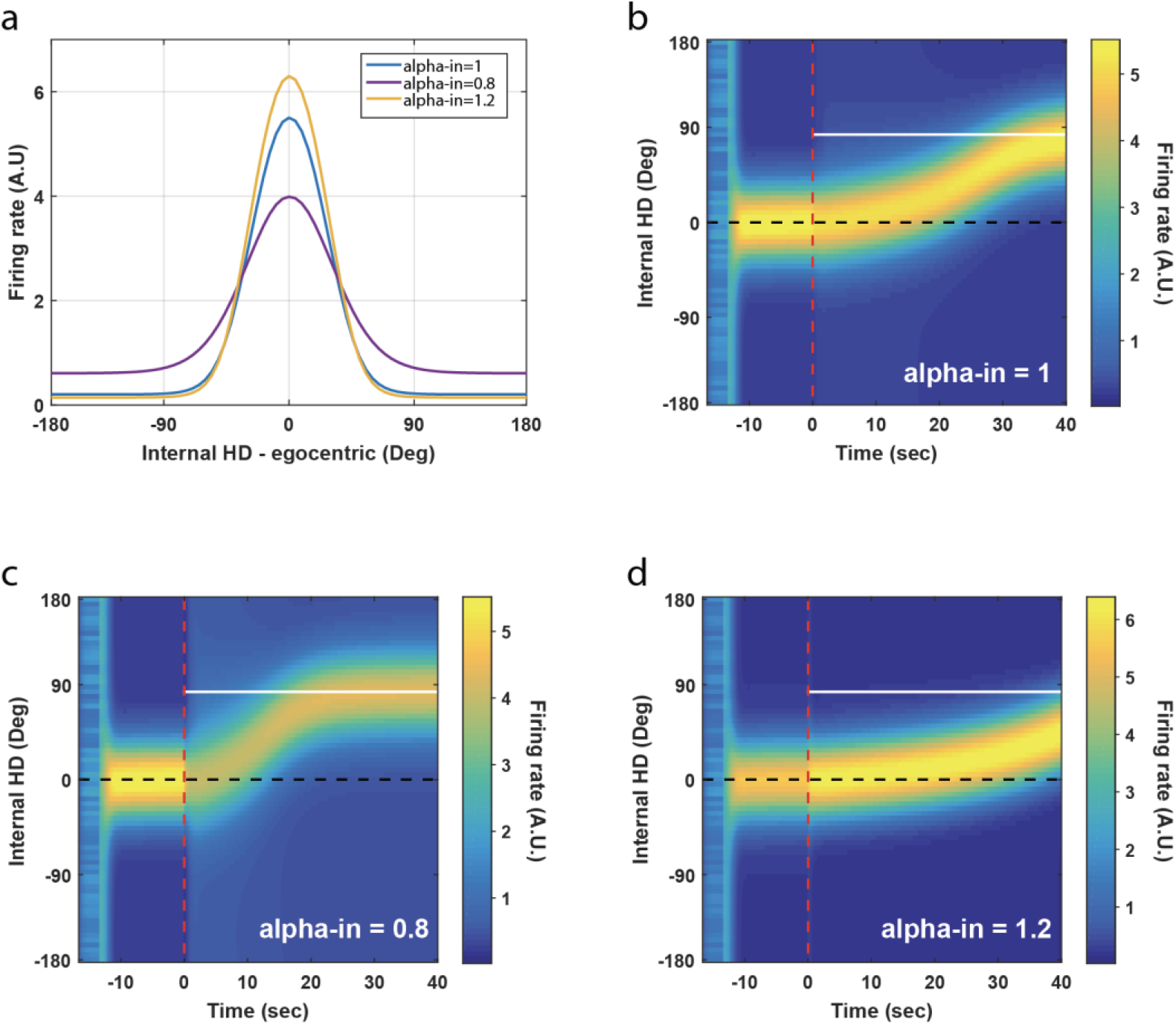
Simulation results showing modulation of reset speed at different gain amplitudes (i.e. “alpha-in”). **a**. Due to weakening lateral inhibition in the HD network, at low gain amplitudes, the egocentric bump of activity exhibits widening of the tuning curve as well as an increase in baseline firing rate. **b, c and d**. Time-dependent dynamics of the bump of activity before and after cue-display (dashed red line). The first 5 seconds ([20:15]s pre-cue) correspond to a random cell activity, after which an external input is applied on the network to impose the formation of the bump of activity at a specific direction (0° in this case). During the pre-cue period, alpha-in is set to 1. Following cue display, alpha-in changes instantaneously and remains at the indicated value for the remainder of the simulation. The 40s post-cue simulation interval may or may not be sufficient to shift and stabilize the network’s HD representation around the visual cue location (solid white line), depending on the gain amplitude. In all these examples, the animal is motionless (Angular velocity = 0°/s).

**Model Table 1:** Simulation parameters

|  |  |  |
| --- | --- | --- |
| $dt$ | Time step | 30ms |
| $N$ | Number of neurons per layer (HD, inhibitory, AV-by-HD and visual layers) | 75 |
| $\tau_i^{HD}$ | Time constant of the decay rate of postsynaptic potential for HD neurons | 300ms |
| $\tau_i^I$ | Time constant of the decay rate of postsynaptic potential for inhibitory neurons | 60ms |
| $\tau_i^{AV-by-HD}$ | Time constant of the decay rate of postsynaptic potential for AV-by-HD neurons | 60ms |
| $\gamma_i^{HD}$ | Tonic inhibition of HD neurons | -1.5 |
| $\gamma_i^I$ | Tonic inhibition of inhibitory neurons | -7.5 |
| $\gamma_i^{AV-by-HD}$ | Tonic inhibition of AV-by-HD neurons | -1 |
| $a$ | Scale parameter of the activation function $F_i$ | 1.1 |
| $b$ | Shift parameter of the activation function $F_i$ | 0.25 |
| $w_{HD \rightarrow I}$ | HD → I connection weights | 3 |
| $w_{HD \rightarrow AV-by-HD}$ | HD → AV-by-HD connection weights | 0.3 |
| $\phi_{offset}$ | Angular offset of AV-by-HD → HD projections | 24° |
| $k_{Vis}$ | Scale parameter of visual input strength | 0.32 |
| $\sigma_{Vis}$ | Standard deviation of the visual input kernel | 11° |
| $k_g$ | Scale parameter of input gain | 4.8 |
| $k_{AV \times HD}$ | Scale parameter of AV-by-HD → HD connection | 1.6248 |
| $\sigma_{AV \times HD}$ | Standard deviation of AV-by-HD → HD weight distribution | 5° |
| $k_I$ | Scale parameter of I → HD connection | 0.0432 |
| $\sigma_I$ | Standard deviation of I → HD weight distribution | 15° |

